# Metabolic compensation via gluconeogenesis explains the non-essentiality of glycogen phosphorylase as an insecticidal target in *Plutella xylostella*

**DOI:** 10.64898/2026.03.03.709364

**Authors:** Yifei Zhou, Yanqi Kang, Yan Liu, Ruichi Li, Dongliang Wang, Chong Yi, Yifan Li, Yalin Zhang, Zhen Tian, Jiyuan Liu

**Affiliations:** Key Laboratory of Plant Protection Resources and Pest Management of Ministry of Education, College of Plant Protection, Northwest A&F University, Yangling, Shaanxi, China; Northwest A&F University Shenzhen Research Institute, Shenzhen, Guangdong, China

**Keywords:** glycogen phosphorylase, benzoylphenylurea, insecticide target, metabolic compensation, gluconeogenesis, Diamondback moth, chitin synthesis

## Abstract

Glycogen phosphorylase (GP) catalyzes the rate-limiting step of glycogenolysis and occupies a central position in insect carbohydrate metabolism, supplying precursors for chitin biosynthesis. Acyl urea compounds structurally related to benzoylphenylurea (BPU) insecticides are potent inhibitors of mammalian GP, raising the question of whether insect GP could serve as an independent insecticidal target. Here, we systematically evaluate this possibility in the diamondback moth *Plutella xylostella*. We show that a mammalian GP inhibitor (GPI) potently inhibits recombinant PxGP (IC_50_ = 2.96 nM), while the BPU insecticide diflubenzuron (DFB) does not. Molecular docking and MM/GBSA analysis predict that this selectivity reflects differential side-chain engagement: GPI is predicted to occupy the allosteric site at the dimer interface via contacts with seven residues spanning both subunits (ΔG = −34.63 kcal/mol), whereas DFB’s difluorobenzoyl moiety is predicted to remain solvent exposed without productive protein contacts (ΔG = −29.29 kcal/mol). Despite potent in vitro inhibition, GPI exhibits no insecticidal activity, and RNAi-mediated knockdown of PxGP (confirmed by enzyme activity measurements showing ∼29% reduction in per-larva GP-a activity) does not impair development or survival. We provide evidence that insects compensate through a multi-layered metabolic response: upregulation of gluconeogenic enzymes (PEPCK, G-6-Pase), selective upregulation of glycogen branching enzyme (GBE) but not α-amylase, and changes in protein content suggestive of catabolic substrate mobilization. Fitness assessment reveals transient larval weight loss (24–48 h) with complete recovery of pupal weight, adult wing morphology, and female fecundity, indicating that the metabolic compensation response is ultimately effective in sustaining development. These findings suggest that GP is functionally non-essential under these conditions, likely through a combination of gluconeogenic compensation and the availability of alternative carbon sources, and highlight the importance of considering metabolic network redundancy in target-based insecticide design.

## Introduction

Benzoylphenylurea (BPU) insecticides, including diflubenzuron (DFB), have been widely used for pest control for over four decades. The molecular basis of BPU-mediated insecticidal activity has been the subject of extensive investigation. Genetic studies, most notably the demonstration through CRISPR/Cas9 gene editing that amino acid substitutions at a single site in chitin synthase (CHS) confer high-level resistance to DFB in *Drosophila melanogaster* [1], provide compelling evidence that CHS is the primary target gene associated with BPU resistance. These findings establish a clear causal genetic link between CHS and BPU activity.

Despite this genetic evidence, the biochemical mechanism by which BPUs inhibit CHS remains incompletely characterized. Multiple in vitro studies of chitin synthase, including those using cell-free chitin synthase preparations, have failed to demonstrate direct enzymatic inhibition by BPU compounds: DFB did not inhibit *Tribolium* gut chitin synthetase [2], and a systematic evaluation concluded that chitin synthesis-inhibiting insect growth regulators do not inhibit chitin synthase in cell-free assays [3, 4]. More recently, Zhang and Zhu (2013) reported only slight in vitro inhibition of *Anopheles gambiae* CHS activity by DFB at the highest concentration tested, with no in vivo inhibition detected [5]. These findings led to the recognition that a key obstacle in identifying the action site of BPU insecticides has been the lack of in vitro demonstration of their ability to inhibit insect chitin synthesis under cell-free conditions. This gap between genetic and biochemical evidence does not diminish the importance of the genetic findings, but it does highlight the need for continued investigation at the enzyme level [6, 7].

Glycogen Phosphorylase (GP) catalyzes the rate-limiting step in glycogenolysis and plays a central role in insect carbohydrate metabolism, providing glucose-1-phosphate that feeds into the hexosamine pathway and ultimately supplies UDP-N-acetylglucosamine for chitin biosynthesis [8, 9]. In insects, GP-derived glucose is converted to trehalose (the primary hemolymph sugar), which is subsequently channeled through a series of enzymatic steps to generate UDP-GlcNAc, the substrate monomer for chitin synthesis. GP also plays key roles in carbohydrate homeostasis, osmotic regulation, stress adaptation [10–13]. Acyl urea compounds structurally related to BPUs have been identified as potent inhibitors of mammalian GP (Figure 1) [14, 15], raising the question of whether insect GP could itself represent a viable insecticidal target — independent of the BPU mechanism-of-action question. However, the feasibility of targeting insect GP for pest control has not been systematically evaluated.

**Figure 1.**
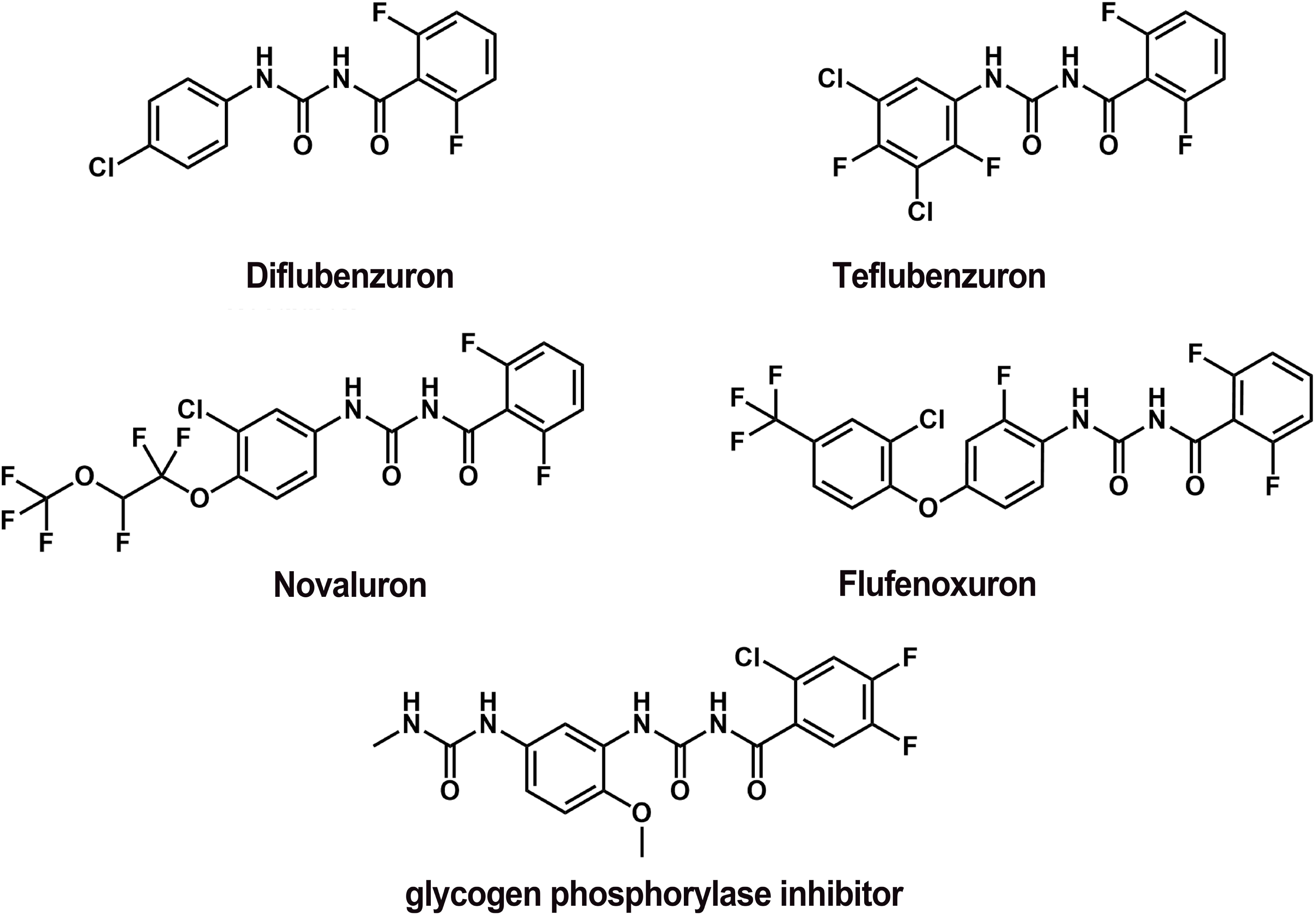
Structural comparison of benzoylphenylurea (BPU) insecticides and a mammalian glycogen phosphorylase inhibitor (GPI) reveals a shared acylurea scaffold. Chemical structures of four BPU insecticides (Diflubenzuron, Teflubenzuron, Novaluron, and Flufenoxuron) and a human glycogen phosphorylase inhibitor (GPI) are shown. All share a central N,N’-diphenylurea (acylurea) core (dashed boxes). BPUs are characterized by halogenated (F, Cl) aromatic substituents and in some cases by additional ether or alkyl side chains. The GPI contains a methoxyphenyl group bearing an additional methylurea side chain.

In this study, we address two distinct questions. First, we test whether DFB directly inhibits *Plutella xylostella* GP (PxGP) at the enzyme level, adding biochemical data to the existing genetic framework. Second, and more importantly, we systematically evaluate GP as an independent insecticidal target by characterizing the metabolic consequences of GP inhibition and knockdown in *P. xylostella*. Our findings reveal that PxGP is functionally non-essential for larval development under the tested conditions, reflecting a previously uncharacterized gluconeogenic compensation mechanism, providing new insight into the metabolic plasticity of insects and its implications for target-based insecticide design.

## Results

### Recombinant *P. xylostella* glycogen phosphorylase (PxGP) is a functional, activatable enzyme

To investigate its biochemical properties and evaluate its potential as an insecticidal target, the full-length *PxGP* cDNA (predicted molecular mass of 98.1 kDa) was cloned and expressed in Sf9 cells using the Bac-to-Bac baculovirus expression system. Following purification via immobilized metal-affinity chromatography (IMAC) of the N-terminal 6×His tagged protein [16], SDS-PAGE analysis and Western blot analysis confirmed a highly pure protein band migrating at approximately 100 kDa, consistent with its theoretical mass (Figure 2A). A total yield of 7.5 mg of purified protein was obtained from 8.25 g of cell pellet (specific yield=0.91 mg/g cells).

**Figure 2.**
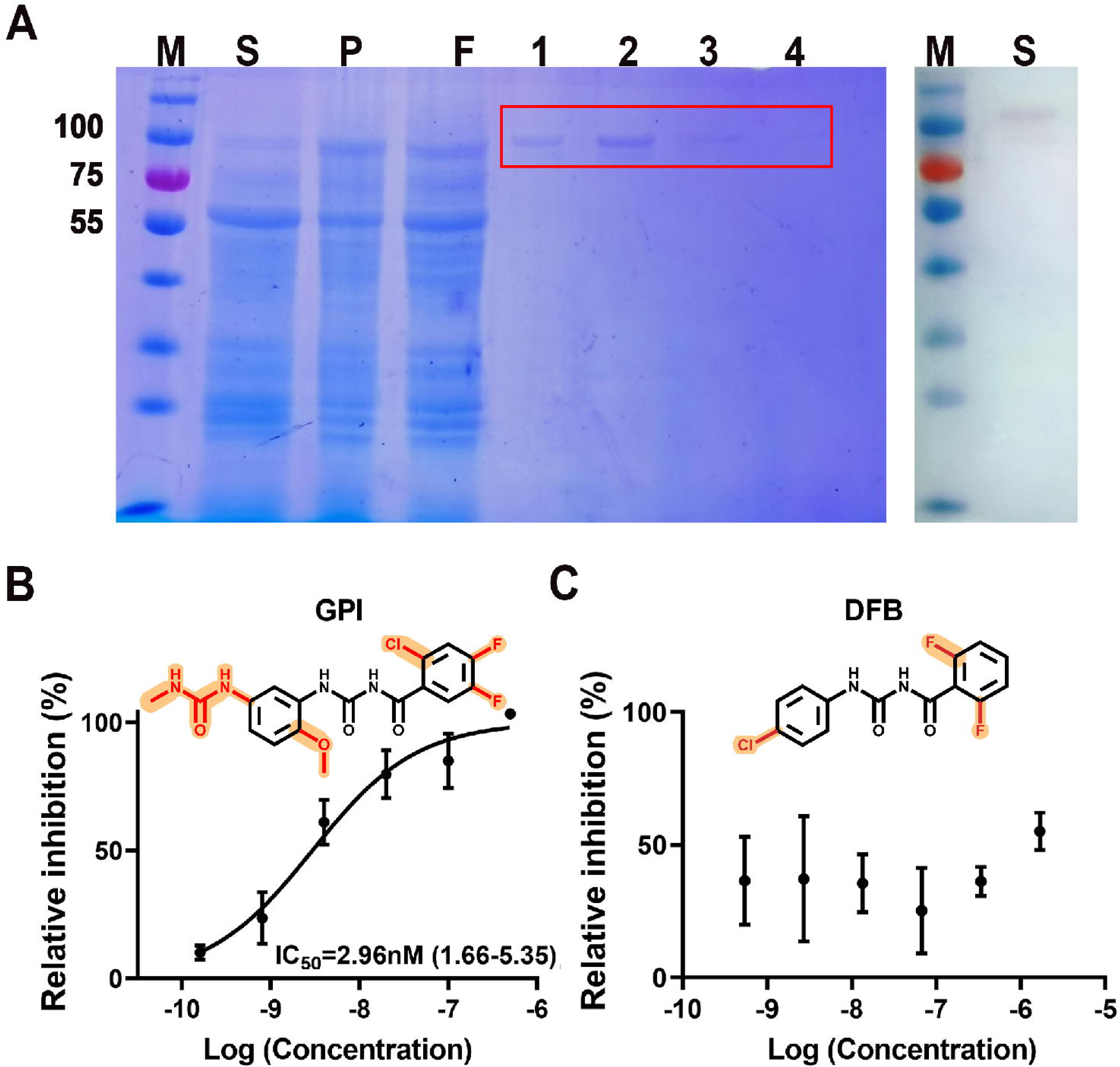
Recombinant PxGP is inhibited by a mammalian GP inhibitor (GPI) but not by the benzoylphenylurea Diflubenzuron. (A) Purification of 6×His-tagged PxGP from Sf9 cells. Left panel: Coomassie-stained SDS-PAGE. Right: Western blot with anti-His antibody. The expected ∼100 kDa band is indicated (red box). M, marker; S, soluble fraction; P, pellet; F, flow-through; lanes 1–4, imidazole elution fractions. (B) Dose-response inhibition by GPI. Activated PxGP-a activity was measured after pre-incubation with GPI (0.16–500 nM). Data (mean ± SEM, n=3) were fit to a four-parameter logistic curve. The calculated IC_50_ is 2.96 nM. *Inset:* GPI structure with its characteristic asymmetric side chains (orange). (C) Inhibition assay with DFB (0.54–1700 nM). DFB did not yield a complete sigmoidal inhibition curve; maximum inhibition did not exceed 55.07 ± 7.05% at the highest concentration. *Inset:* DFB structure with its characteristic halogenated benzoyl groups (orange). This rules out PxGP as a direct target of DFB.

Glycogen phosphorylase activity is regulated by reversible phosphorylation, existing as a less-active dephosphorylated b form (PxGP-b) and a fully active phosphorylated a form (PxGP-a) [17]. Recombinant PxGP-b was activated by incubation with immobilized phosphorylase kinase (PhK) in the presence of ATP and Mg^2+^ for 2 h [18]. The activated enzyme showed robust catalytic activity in a coupled-enzyme spectrophotometric assay, with NADPH production (monitored by increase in A_340_) increasing linearly for the first 15 min and reaching a plateau by approximately 20 min, consistent with substrate depletion and/or product accumulation. The specific activity of activated PxGP-a was 294.25 ± 3.45 U/mg protein (Figure 2–figure supplement 1A), confirming that the recombinant enzyme is catalytically competent for inhibitor screening.

### GPI potently inhibits PxGP *in vitro* and *in vivo*

Given the structural similarity between insecticidal BPUs and mammalian GPIs [15] (Figure 1), we tested whether PxGP could be a target of acylurea-scaffold compounds. GPI exhibited potent, dose-dependent inhibition of recombinant PxGP-a, with a sigmoidal dose-response curve typical of competitive or allosteric inhibition (Figure 2B). Non-linear regression analysis yielded an IC_50_ value of 2.96 nM (95% confidence interval [CI]: 1.66–5.35 nM, Hill slope = 0.74, R² = 0.877, n = 3). Near-complete inhibition (>95%) was achieved at 500 nM GPI, indicating saturable, high-affinity binding. The potency of GPI against PxGP is comparable to its effect on mammalian orthologs (see below) [15].

In striking contrast, the BPU insecticide Diflubenzuron (DFB), which shares the acylurea core with GPI (Figure 1), did not effectively inhibit PxGP-a. No standard sigmoidal dose-response curve was observed over 0.54 nM to 1700 nM (Figure 2C). Maximum inhibition observed did not exceed 55.07 ± 7.05% (mean ± SEM), with high variability at lower concentrations. These data indicate that DFB, unlike GPI, is not an effective inhibitor of PxGP, ruling out GP as the direct molecular target of this BPU insecticide.

Consistent with the recombinant enzyme results, GPI effectively inhibited native GP activity in crude lysates from third-instar larvae in a dose- and time-dependent manner (Figure 2–figure supplement 1B). After 45 min of pre-incubation, 10 µM GPI reduced native GP activity by 57.52 ± 1.88% (n = 3). DFB, however, showed no significant inhibitory effect on native GP activity at any concentration tested (10^-7^, 10^-6^, or 10^-5^ M) or time point (5, 10, 20, or 30 min), with activity remaining at 97-108% of control (Figure 2–figure supplement 1C).

Collectively, these results establish PxGP as a high-affinity molecular target for GPI, whereas DFB does not inhibit PxGP.

### Paradoxical finding: potent PxGP inhibition by GPI lacks *in vivo* toxicity

Because GPI potently inhibits PxGP, the rate-limiting enzyme in glycogenolysis, which supplies precursors for chitin synthesis [19], we expected it to exhibit significant insecticidal activity. In leaf-dip bioassays with third-instar *P. xylostella* larvae continuously exposed to GPI (250 or 500 mg/L) or DFB (125 or 250 mg/L), GPI caused no significant toxicity or developmental disruption across the entire time course (24 to 120 h post-treatment) (Figure 3A). At 120 h, mortality with 500 mg/L GPI was only 12.78 ± 2.42% (n = 3), which was not significantly different from the solvent control (CK: 5.00 ± 5.00%, *P* = 0.21) (Figure 3B). Larvae exposed to 250 mg/L GPI exhibited even lower mortality (2.50 ± 2.50%).

**Figure 3.**
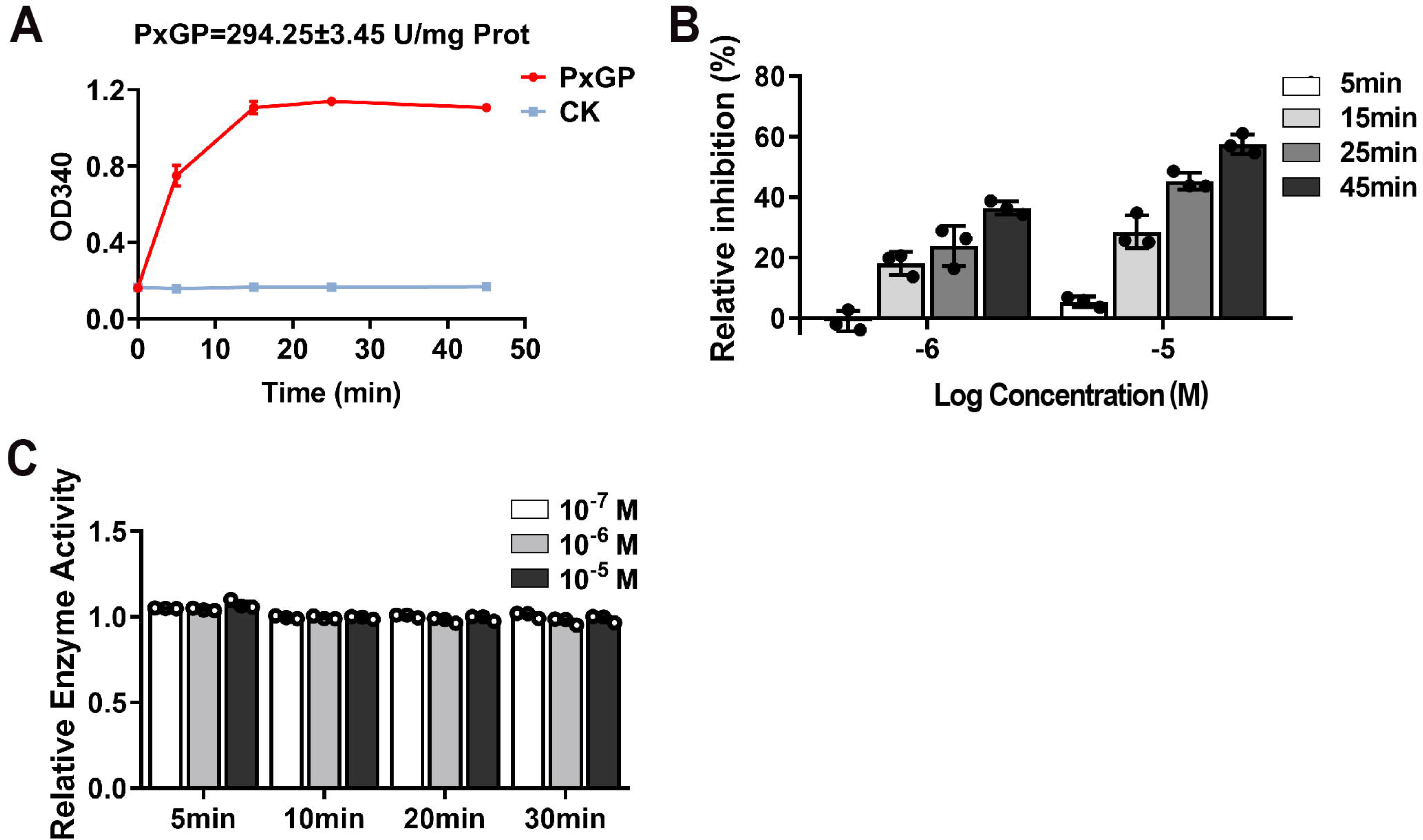
The GP inhibitor (GPI) is non-toxic to *Plutella xylostella*, unlike the benzoylphenylurea Diflubenzuron (DFB). (A) Developmental stage distribution over time for control, GPI (250, 500 mg/L), and DFB (125, 250 mg/L) treated larvae. DFB causes developmental arrest. (B) Quantitative effects on mortality (top), pupation (middle), and adult emergence (bottom). GPI groups did not differ from control (*P* > 0.05), whereas DFB caused significant, dose-dependent mortality and severely impaired development (*P* < 0.05). Data are mean ± SEM (n=3). Different letters denote statistical significance. (C) Larval phenotypes at 96 h. Control and GPI-treated larvae are normal. DFB-treated larvae show catastrophic molting failure and “double cuticle” (red arrows). Scale bar, 1 mm. (D) Pupal and adult phenotypes. GPI-treated individuals develop normally. DFB treatment results in deformed pupae and trapped pharate adults (red arrows). Scale bar, 1 mm.

In stark contrast, DFB induced significant, dose-dependent mortality. At 120 h, DFB treatment at 125 mg/L caused 41.60 ± 4.85% mortality (*P* < 0.01 vs. control) and at 250 mg/L caused 61.36 ± 10.02% mortality (*P* < 0.01 vs. control). Probit analysis of the DFB dose-response data yielded calculated LC_30_ and LC_50_ values of 141.900 mg/L (95% CI: 67.893–206.576 mg/L) and 229.537 mg/L (95% CI: 145.287-331.784 mg/L), respectively, with an acceptable goodness-of-fit (χ² = 11.946, *P* = 0.683) (Supplementary file 2).

Furthermore, GPI-treated larvae showed no developmental abnormalities. Pupation rates were 64.17 ± 4.79% for the 500 mg/L GPI group and 69.72 ± 6.81% for the 250 mg/L group, which were statistically indistinguishable from the control group (64.72 ± 9.33%, *P* > 0.05 for both comparisons) (Figure 3B, middle panel). Similarly, adult emergence rates were 56.11 ± 5.64% and 56.39 ± 6.24% for the 500 and 250 mg/L GPI groups, respectively, compared to 54.10 ± 11.91 % for the control (*P* > 0.05, Figure 3B). Phenotypically, GPI-treated larvae molted normally, fed actively, and displayed no cuticle defects (Figure 3C). DFB-treated larvae exhibited the characteristic phenotypes of BPU intoxication: catastrophic molting failure, larval-pupal intermediates, and complete failure of adult emergence (Figure 3C and 3D). This striking discrepancy between biochemical potency and *in vivo* inefficiency presents a profound paradox that demands mechanistic investigation.

### GPI exposure triggers a compensatory upregulation of *PxGP* transcription

To explore this discrepancy, we measured *PxGP* gene expression after GPI exposure. RT-qPCR revealed that 500 mg/L GPI rapidly upregulated *PxGP* mRNA transcription. As shown in Figure 4, its relative expression increased 3.24-fold at 24 h, peaked at 3.48-fold at 48 h, and remained elevated at 96 h. This indicates that the insect responds to the pharmacological inhibition by transcriptionally increasing the *de novo* synthesis of the target enzyme.

**Figure 4.**
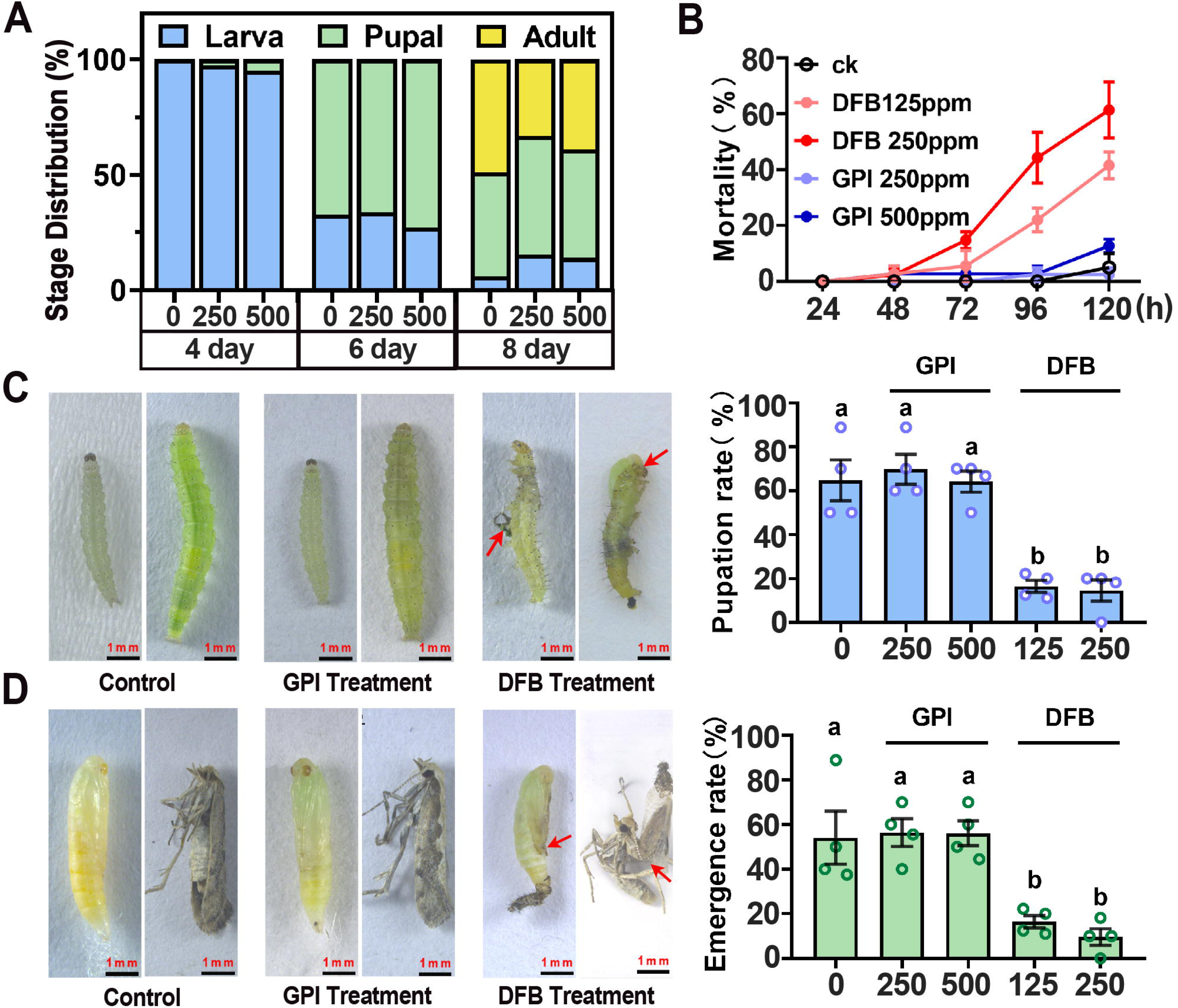
Ingestion of GP inhibitor (GPI) induces compensatory upregulation of ***PxGP* transcription.** Relative *PxGP* mRNA levels in larvae after continuous exposure to 500 mg/L GPI or control (CK). Expression was normalized to *PxRPS13* and calibrated to the 24 h control sample (set to 1.0) using the 2^-ΔΔCt^ method. Data are mean ± SEM (n=3). GPI triggered a rapid and sustained transcriptional increase, peaking at 48 h (3.48-fold). Significance vs. control: \**P* < 0.05, \*\**P* < 0.01, \*\*\**P* < 0.001 (two-way ANOVA with Sidak’s test).

### Sequence homology and rationale for target site selection

To achieve effective gene knockdown coupled with functional suppression, precise selection of the interference target site is critical. Sequence analysis of PxGP (837 residues) revealed that residues 88-680 form the active site pocket of the GT35_Glycogen_Phosphorylase family, and the C-terminal region contains a phosphorylase pyridoxal-phosphate attachment site (residues 672-684). Eleven residues within positions 42-318 are conserved and correspond to the AMP-binding site in human GP (Figure 5A) [20].

**Figure 5.**
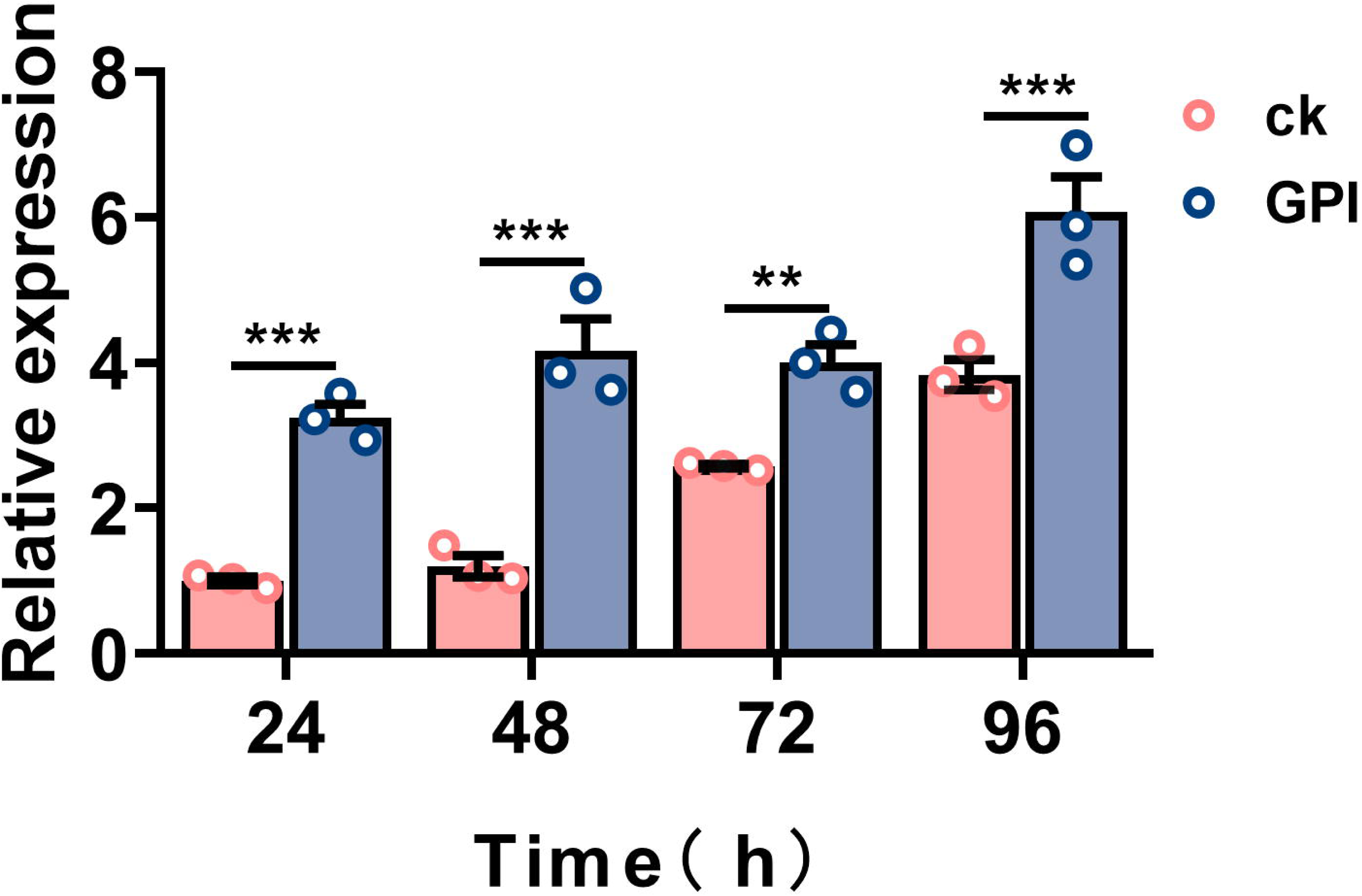
PxGP sequence conservation and domain architecture. (A) Schematic of PxGP functional domains: pyridoxal phosphate (PLP) binding site (blue boxes), AMP allosteric site (yellow boxes), and glycogen-binding/active site pocket (blue shading). (B) Multiple sequence alignment of GP from *Plutella xylostella* (PxGP), other lepidopterans (*Helicoverpa armigera* HaGP, *Spodoptera exigua* SeGP, *Ostrinia furnacalis* OfGP), human (HsGP), and rabbit (OcGP). Identical residues are shaded red. The gray box indicates the 606 bp region (aa 22–224) targeted for RNAi, which overlaps the conserved AMP-binding domain. PxGP shares 82.29% similarity with human GP.

Multiple sequence alignment of PxGP with GP orthologs from other key lepidopteran pests (*Helicoverpa armigera* HaGP, *Spodoptera exigua* SeGP, *Ostrinia furnacalis* OfGP), human (HsGP), and rabbit (OcGP) demonstrated high conservation across species (Figure 5B). PxGP shares 82.29% sequence similarity with human GP (Figure 5–figure supplement 1). Since GP activity is allosterically regulated by AMP binding and phosphorylation at Ser14, triggering a conformational shift from the inactive T-state GPb (unphosphorylated) to the active R-state GP-a (phosphorylated) [17], we targeted the core AMP-binding site region (gray box in Figure 5B) for RNAi. Theoretically, interference in this region should prevent AMP-mediated activation of existing GPb while degrading the mRNA pool, substantially reducing de novo GP protein synthesis and maximizing functional suppression.

### RNAi confirms PxGP is non-essential for larval development

The compensatory upregulation of PxGP mRNA expression in response to pharmacological inhibition (GPI exposure experiments) indicated that the gene is under active transcriptional regulation and suggested that increased enzyme synthesis might partially mitigate the effects of GPI. However, this compensation alone could not fully explain the complete absence of phenotype. To determine whether a more profound loss-of-function—achieved by directly suppressing *PxGP* gene expression—is lethal, we performed RNAi to knock down *PxGP* at the transcriptional level.

Firstly, we quantified the expression profile of *PxGP* across developmental stages in *P. xylostella*, *PxGP* was highly expressed in fourth-instar larvae (L4) and adults (Figure 6), stages that correlate with the energy demands for metamorphosis and flight, respectively [8, 21]. Given the inherent time lag required for systemic dsRNA delivery to achieve effective post-transcriptional knockdown [22], dsRNA was administered during the mid-third-instar larval stage. Mid-third-instar larvae were microinjected with dsRNA targeting a conserved 606 bp region of *PxGP* (ds*GP*) or with a non-specific control ds*GFP*. Three concentrations of ds*GP* (6,000, 10,000, and 14,000 ng/µL) were tested, and knockdown efficiency was assessed by RT-qPCR at 24, 48, 72, and 96 h post-injection (Figure 7). The highest concentration (14,000 ng/µL) achieved maximal suppression at 48 h, reducing *PxGP* transcript levels to 12.47 ± 1.68% of the ds*GFP* control (87.59% knockdown, *P* < 0.001) (Figure 7B).

**Figure 6.**
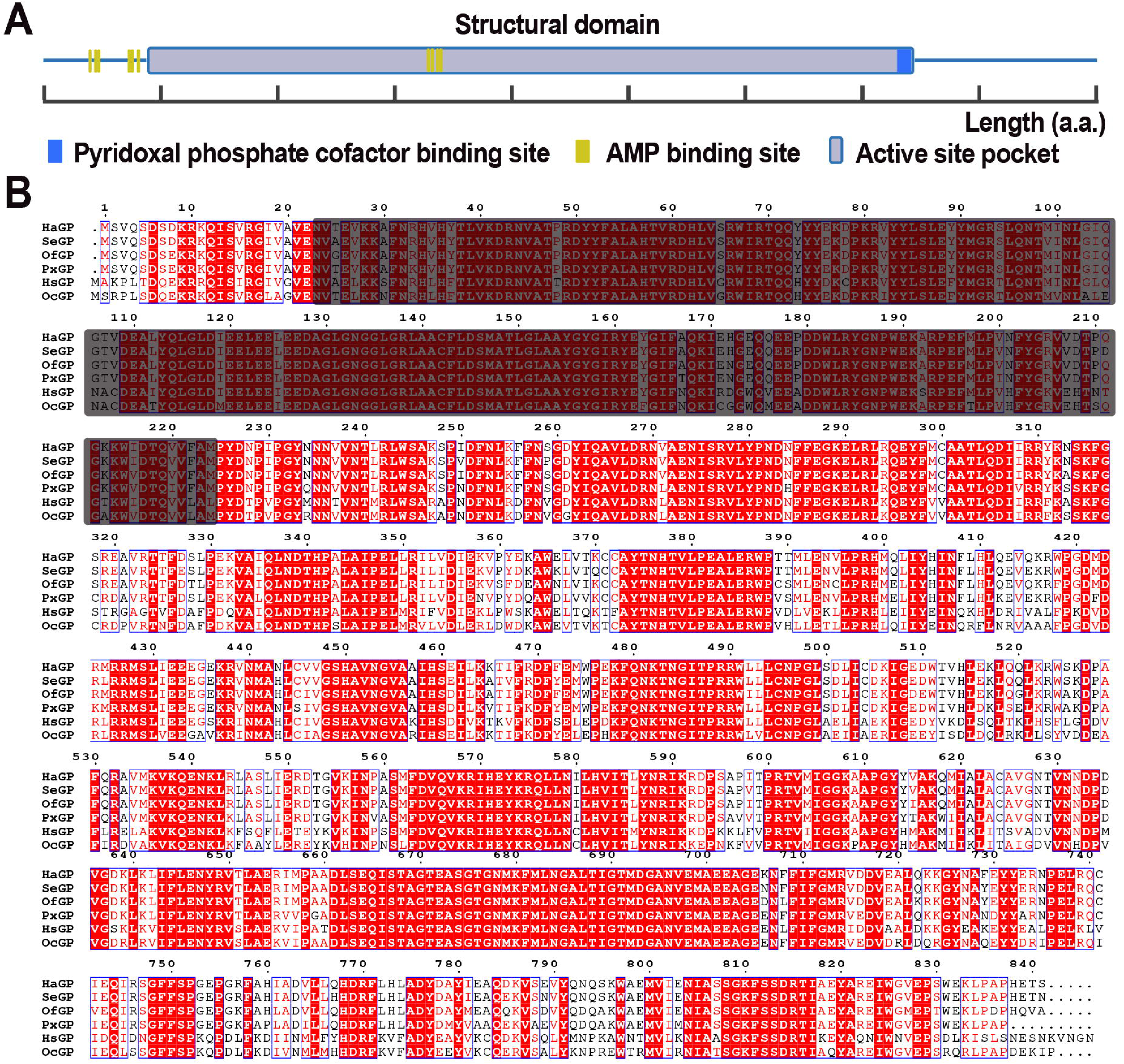
Developmental expression profile of *PxGP*. Relative *PxGP* mRNA levels across life stages (Egg, larval instars L1-L4, Pupa, Adult), normalized to *PxRPS13* (Egg set to 1.0). Data are mean ± SEM (n=3). Expression peaks in adults. Different letters indicate significant differences (*P* < 0.05, one-way ANOVA with Tukey’s test).

**Figure 7.**
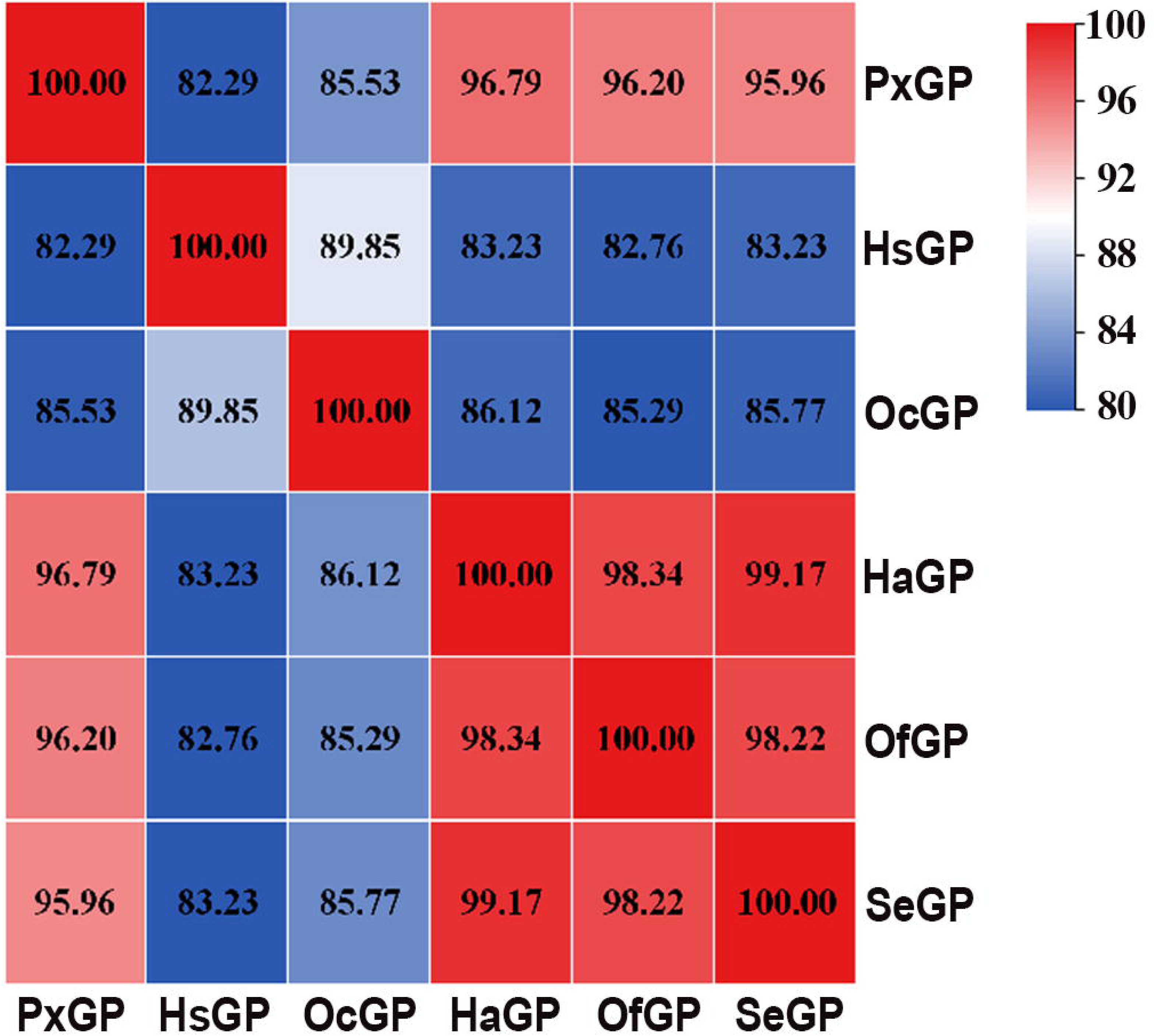
RNAi-mediated knockdown of *PxGP* is dose- and time-dependent. (A) Experimental workflow. Third-instar larvae were injected with dsRNA targeting *PxGP* (ds*G*P) at three concentrations or a control dsRNA (ds*GFP*), then sampled for analysis. (B) Time-course of *PxGP* mRNA levels post-injection, relative to ds*GFP* control (set to 1.0). The highest ds*GP* dose (14,000 ng/µL) caused maximal suppression (87.6%) at 48 h. Data are mean ± SEM (n=3). Significance vs. ds*GFP* at each time point: * *P* < 0.05, \*\**P* < 0.01, \*\*\**P* < 0.001 (one-way ANOVA with Tukey’s test).

Significant knockdown persisted at 72 h (69.86% suppression, *P* < 0.05), with partial recovery by 96 h, likely due to dsRNA degradation and/or compensatory transcriptional upregulation. The 10,000 ng/µL dose also produced significant knockdown at 48 h (65.76% suppression, *P* < 0.001), whereas the 6,000 ng/µL dose showed more modest effects (Figure 7B).

Despite this robust, transient silencing, no adverse phenotypic effects were observed. Developmental stage distribution and mortality rates in all ds*GP*-treated groups (6,000, 10,000, and 14,000 ng/µL) were statistically indistinguishable from the ds*GFP* control group across the entire 120 h observation period (*P* > 0.05 at all time points) (Figure 8A and 8B). At 120 h, cumulative mortality was 8.10 ± 4.23%, 9.82 ± 0.81%, and 3.33 ± 3.33% for the 6,000, 10,000, and 14,000 ng/µL ds*GP* groups, respectively, comparable to the ds*GFP* control (Figure 8B). Pupation and adult emergence were also indistinguishable from controls (Figure 8B). Phenotypic examination revealed that ds*GP*-injected larvae developed normally, molted successfully, formed morphologically normal pupae, and emerged as viable adults without cuticle defects or developmental delays (Figure 8C and 8D). Thus, acute, profound suppression of *PxGP* expression is not lethal to *P. xylostella* during the critical larval-pupal transition, despite the enzyme’s established role in glycogenolysis and chitin precursor synthesis. This finding necessitates an investigation into the compensatory mechanisms that allow the insect to survive in the absence of functional GP.

**Figure 8.**
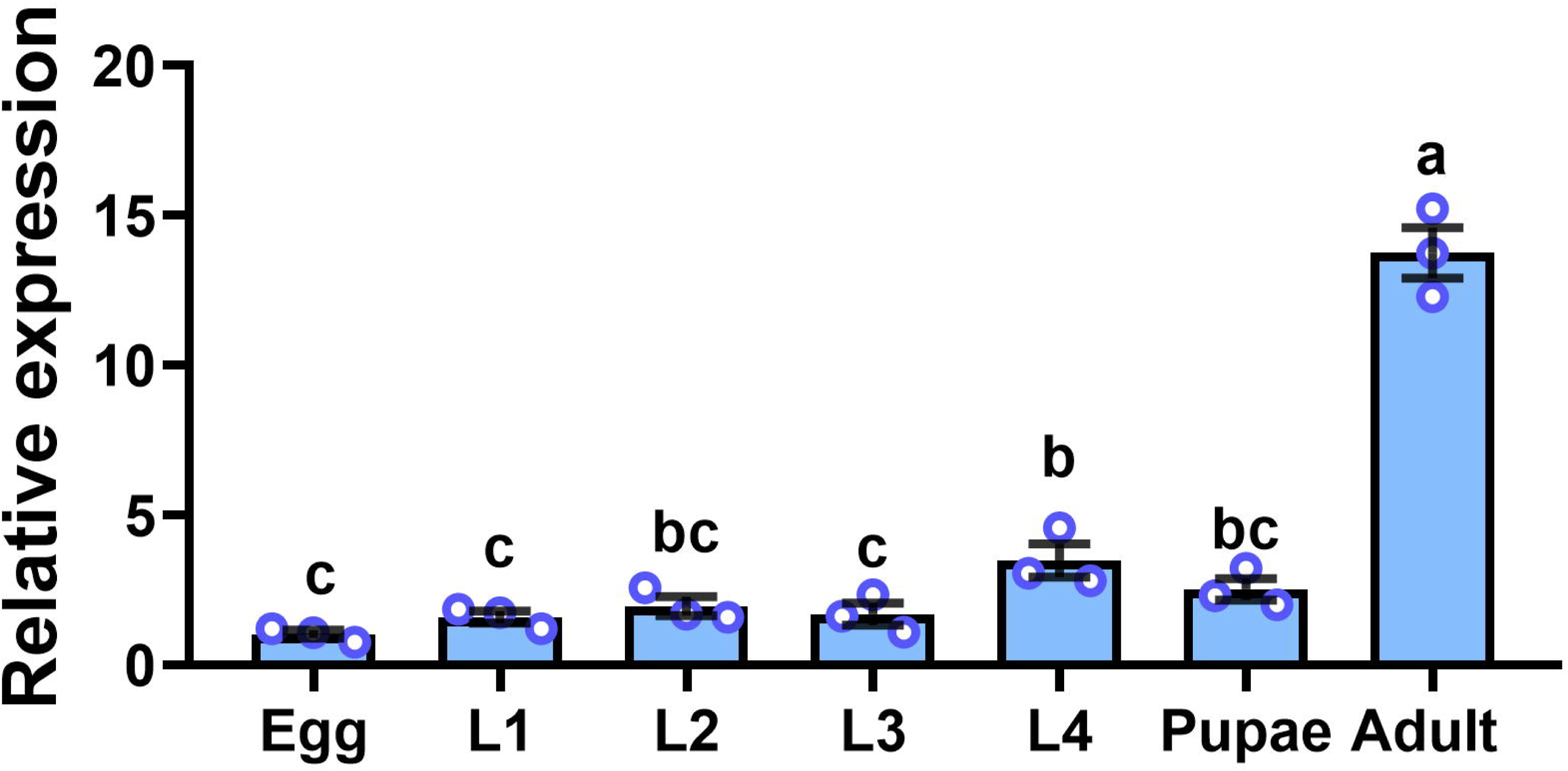
*PxGP* knockdown does not impair development or survival. (A) Developmental stage distribution post-injection shows similar progression across all groups. (B) Cumulative mortality (top), pupation (middle), and adult emergence (bottom) rates were not significantly different between ds*GP* and ds*GFP* control groups (*P* > 0.05). Data are mean ± SEM (n=3). (C) Representative larval phenotypes at 48 h and 96 h post-injection are normal in both ds*GFP* and high-dose ds*GP* groups. Scale bar, 1 mm. (D) Pupae and adults from both groups are morphologically normal. Scale bar, 1 mm.

### *PxGP* knockdown coordinately suppresses the glycogenolysis-to-chitin pathway

To elucidate how larvae survive GP loss, we examined downstream genes linking glycogenolysis to chitin synthesis. Specifically, we measured the expression of trehalase (*PxTre*), which hydrolyzes circulating trehalose to glucose [8], and hexokinase (*PxHex*), which phosphorylates glucose to glucose-6-phosphate—a key branch point leading to UDP-GlcNAc synthesis for chitin production [23, 24].

RT-qPCR analysis revealed that suppression of *PxGP* triggered a coordinated transcriptional downregulation of both downstream genes (Figure 9A and 9B). At 48 h post-injection, coinciding with the peak of *PxGP* knockdown (87.59% suppression), *PxTre* expression was reduced to 30.46 ± 1.25% of the ds*GFP* control (69.54% suppression, *P* < 0.001), and *PxHex* to 22.82 ± 3.39% of control (77.18% suppression, *P* < 0.001). This suppression persisted at 72 h (*PxTre*: 61.27 ± 2.21% suppression, *P* < 0.001; *PxHex*: 41.96 ±4.02% suppression, *P* < 0.001) and by 96 h had returned to, and modestly exceeded, control levels, mirroring the recovery kinetics of *PxGP* itself.

**Figure 9.**
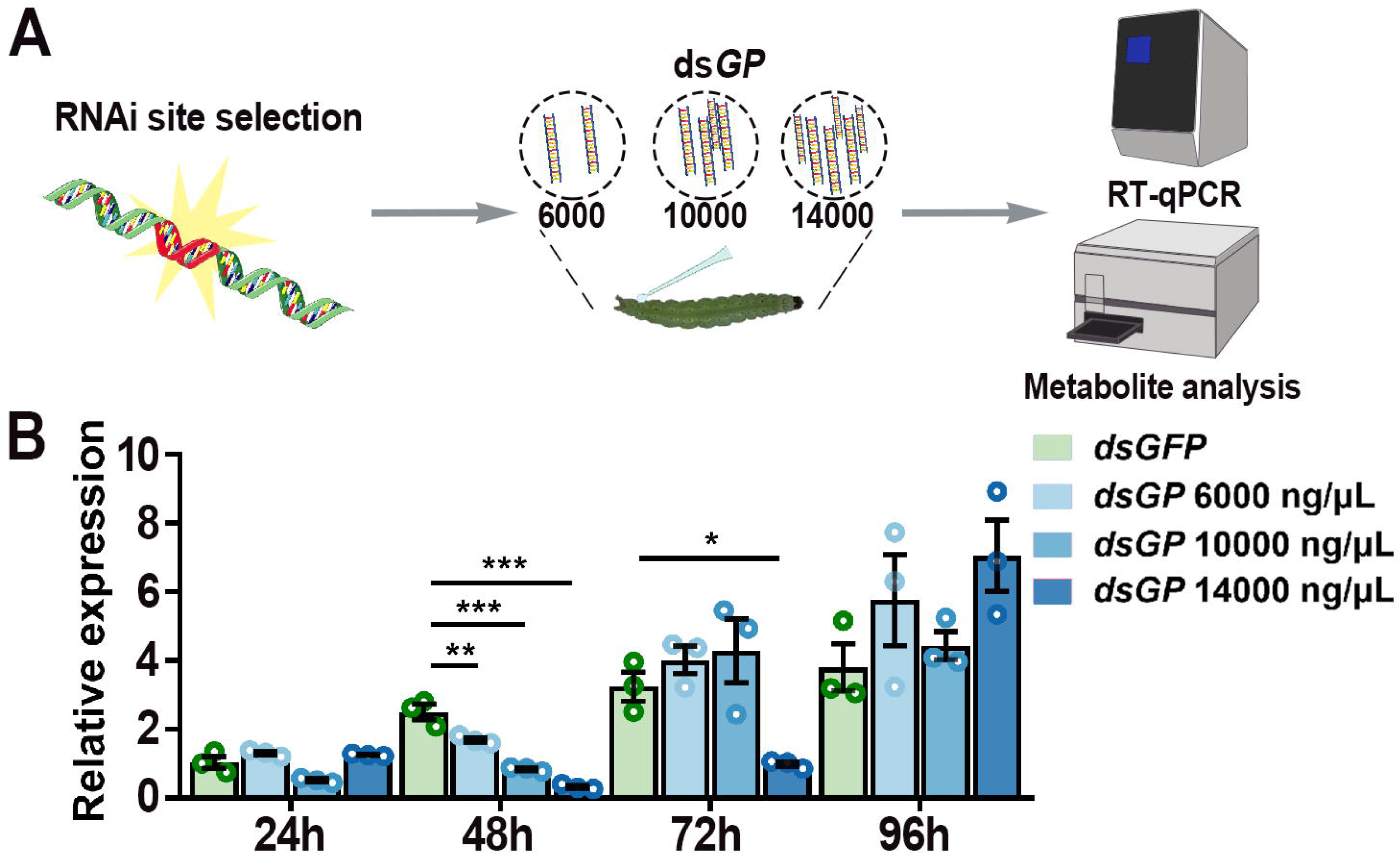
*PxGP* knockdown triggers biphasic metabolic reprogramming, culmina ting in gluconeogenic compensation. (A, B) Concurrent suppression of downstream genes *Trehalase (PxTre)* and *Hexokinase (PxHex)* post-knockdown. (C) Upregulation of gluconeogenic genes *PxPEPCK* and *PxG*-*6*-*Pase* at 96 h. Data in A-C are mean ± S EM (n=3); \**P* < 0.05, \*\**P* < 0.01, \*\*\**P* < 0.001 vs. ds*GFP* control. (D-H) Metabolite levels post-knockdown. (D) Total protein decreased at 72 h (*P* < 0.05). (E) Glycogen remained stable. (F) Trehalose levels were elevated 7.4 fold relative to control at 96 h (*P* < 0.05). (G) Glucose-6-phosphate (G6P) increased 6.5-fold at 96 h (*P* < 0.0 5). (H) Free glucose showed a transient dip at 72 h (*P* < 0.05) before recovery. Data ar e mean ± SEM (n=3); \**P* < 0.05 vs. control at each time point.

This coordinated downregulation indicates that the insect responds to the loss of GP not by futilely attempting to maintain flux through a pathway that lacks its key upstream enzyme, but rather by implementing a broader transcriptional shutdown of the glycogenolysis-to-glucose-to-chitin pathway. This suggests a coherent metabolic response program, raising the question: if glycogenolysis is blocked, how does the insect obtain the glucose required for survival and chitin synthesis?

### Biphasic metabolic response: transient depletion followed by gluconeogenic rescue

The absence of a lethal or developmental phenotype despite glycogenolysis pathway suppression suggested activation of alternative glucose synthesis pathway to compensate for GP loss. We therefore measured expression of key gluconeogenic enzymes and metabolite levels across the glycogeno-glucose axis.

### Gene expression analysis is consistent with gluconeogenic activation

We assessed transcript levels of two rate-limiting gluconeogenic enzymes: phosphoenolpyruvate carboxykinase (PEPCK, encoded by *PxPEPCK*), which catalyzes the conversion of oxaloacetate to phosphoenolpyruvate; glucose-6-phosphatase (G-6-Pase, encoded by *PxG-6-Pase*), which catalyzes the dephosphorylation of glucose-6-phosphate to free glucose. These enzymes are reciprocally regulated relative to glycolysis/glycogenolysis and serve as molecular markers of gluconeogenic flux [25, 26].

At 96 h post-injection―a critical pre-pupal stage with high carbohydrate demand for pupal cuticle synthesis [27, 28]―both *PxPEPCK* and *PxG-6-Pase* were significantly upregulated in ds*GP*-treated larvae (3.95 ± 0.44-fold and 3.34 ± 0.20-fold versus ds*GFP* control, respectively; *P* < 0.001 for each) (Figure 9C). This indicates robust transcriptional activation of the gluconeogenesis pathway in response to GP loss.

### Changes in protein content suggestive of catabolic substrate mobilization

To identify the carbon source for gluconeogenesis, we quantified total protein content in ds*GP*-treated larvae across the time course. Strikingly, a significant 29.39% reduction in protein was observed at 72 h post-injection (*P* < 0.05), coinciding precisely with the nadir of glucose insufficiency (Figure 9D).

This protein decline is consistent with substrate mobilization: the degradation of cellular proteins liberates amino acids, which serve as the primary carbon source for gluconeogenesis in insects [25, 29–31]. The magnitude of protein loss (∼30%) is sufficient to provide substantial amino acid flux into the gluconeogenic pathway.

### Metabolite profiling reveals a biphasic adaptation

Quantification of four key metabolites (glycogen, trehalose, glucose-6-phosphat [G6P], and free glucose) along the glycogen-glucose axis at 48, 72, and 96 h post-injection uncovered a dynamic, biphasic response (Figure 9E-9H).

1. Glycogen Content: Contrary to the expectation of accumulation upon GP blockade, glycogen levels remained unchanged relative to controls throughout the time course (*P* > 0.05) (Figure 9E). This paradox suggests that insects activate alternative, GP-independent pathways for glycogen catabolism―most likely via α-amylase or glycogen debranching enzymes [32–34]―to provide emergency glucose mobilization during the period of metabolic stress, suggesting that these compensatory pathways are sufficient for short-term energy needs but are eventually superseded by gluconeogenic glucose production.
2. Trehalose: Trehalose levels remained stable at 48-72 h but increased substantially by 96 h (7.44-fold elevation, *P* < 0.05) (Figure 9F). This increase coincided with the significant upregulation of gluconeogenic genes (e.g., *PEPCK* and *G-6-Pase*), consistent with the idea that gluconeogenesis contributes to trehalose synthesis and ensures adequate carbohydrate reserves for the upcoming pupation [35]. However, it should be noted that the large fold change at 96 h primarily reflects a pronounced decline in trehalose levels in the ds*GFP* control group at this time point (Figure 9F), rather than a net increase in absolute trehalose content in ds*GP* treated larvae. Thus, rather than indicating accumulation above the initial baseline, the data suggest that GP suppressed larvae maintain their trehalose pool at a time when controls are actively depleting it—a pattern consistent with sustained gluconeogenic input.
3. G6P: G6P, which sits at the intersection of glycolysis, gluconeogenesis, the pentose phosphate pathway, and glycogen synthesis [36], exhibited a pattern mirroring that of trehalose (Figure 9G). G6P content declined gradually from 48 to 72 h, consistent with reduced glucose mobilization from glycogen, but then diverged at 96 h, with ds*GP*-treated larvae maintaining higher levels (6.49-fold elevation, *P* < 0.05). This relative elevation is consistent with the pattern observed for trehalose. Together, the coordinated behavior of G6P and trehalose is consistent with gluconeogenesis-derived glucose being converted into storage and transport carbohydrates.
4. Free Glucose: Free glucose levels showed the most subtle but perhaps most physiologically informative pattern (Figure 9H). At 48 h―coinciding with peak PxGP suppression―glucose content was statistically indistinguishable from controls (*P* > 0.05), indicating that initial homeostatic mechanisms (including GP-independent glycogen breakdown and dietary carbohydrate absorption) successfully buffered against acute GP loss. However, at 72 h, the nadir of the metabolic stress response, free glucose levels declined significantly, albeit modestly (18.15% reduction, *P* < 0.05). This transient dip reveals a critical window of metabolic insufficiency during which the insect’s compensatory mechanisms are insufficient to fully maintain glucose homeostasis. Importantly, by 96 h, glucose levels fully recovered to control values (*P* > 0.05), demonstrating that the gluconeogenic pathway successfully restores glucose homeostasis before critical developmental transitions occur.

### Integrated interpretation

Collectively, these metabolite and gene expression data reveal a biphasic metabolic adaptation that provides a coherent explanation for why substantial GP suppression causes no mortality or developmental defects:

1. Phase I (emergency compensation, 48-72 h): Metabolic stress and emergency compensation during the initial response to PxGP suppression, the insect experiences genuine metabolic stress, evidenced by the glycogen levels neither accumulated nor were depleted and the transient reduction in free glucose at 72 h. The static glycogen content indicates activation of GP-independent catabolism pathways, which provide emergency glucose but are insufficient for complete homeostatic maintenance. The modest glucose dip at 72 h demonstrates that this stress is real but tolerable, likely due to several factors: (i) the magnitude is modest (only 18.15% reduction), (ii) the duration is brief (∼24 h), (iii) the timing coincides with the mid-L4 larval stage when energy demands are moderate and active feeding allows dietary carbohydrate supplementation, and (iv) trehalose stores (which remain stable during this phase) can be mobilized to buffer hemolymph glucose.
2. Phase II (gluconeogenic rescue, 96 h): Robust transcriptional upregulation of *PEPCK* and *G-6-Pase* (3-4-fold increases) was observed under GP suppression, which is consistent with enhanced gluconeogenic capacity. At 96 h, trehalose and G6P levels in ds*GP* treated larvae were maintained at higher levels than those in the ds*GFP* controls, which had declined markedly by this time point. This relative maintenance suggests that larvae under GP suppression sustain their carbohydrate pools at a stage when control larvae are actively depleting them. Such maintenance may help ensure adequate carbohydrate supply for the energetically demanding process of pupal cuticle synthesis, which begins shortly after 96 h.

This substrate-to-product chain for gluconeogenesis-mediated metabolic rescue— protein degradation → amino acid release → gluconeogenic conversion → glucose synthesis → trehalose storage—constitutes a complete compensatory mechanism that sustains development despite substantial GP suppression.

### RNAi-mediated suppression of *PxGP* reduces enzyme activity *in vivo*

To confirm that RNAi-mediated transcript suppression of PxGP translates to reduced enzyme function *in vivo*, we measured glycogen phosphorylase a (GP-a) activity in crude extracts from ds*GP*-treated and ds*GFP*-treated larvae at 24, 48, 72, and 96 h post-injection using a coupled-enzyme spectrophotometric assay (Figure 10A, B). At 48 h post-injection in this cohort, *PxGP* mRNA levels were reduced by 54.66% (Figure 10–figure supplement 1). This is lower than the 87.59% suppression achieved in the dose–response experiment (Figure 7B), reflecting batch-to-batch variation in RNAi efficiency between independently injected cohorts.

**Figure 10.**
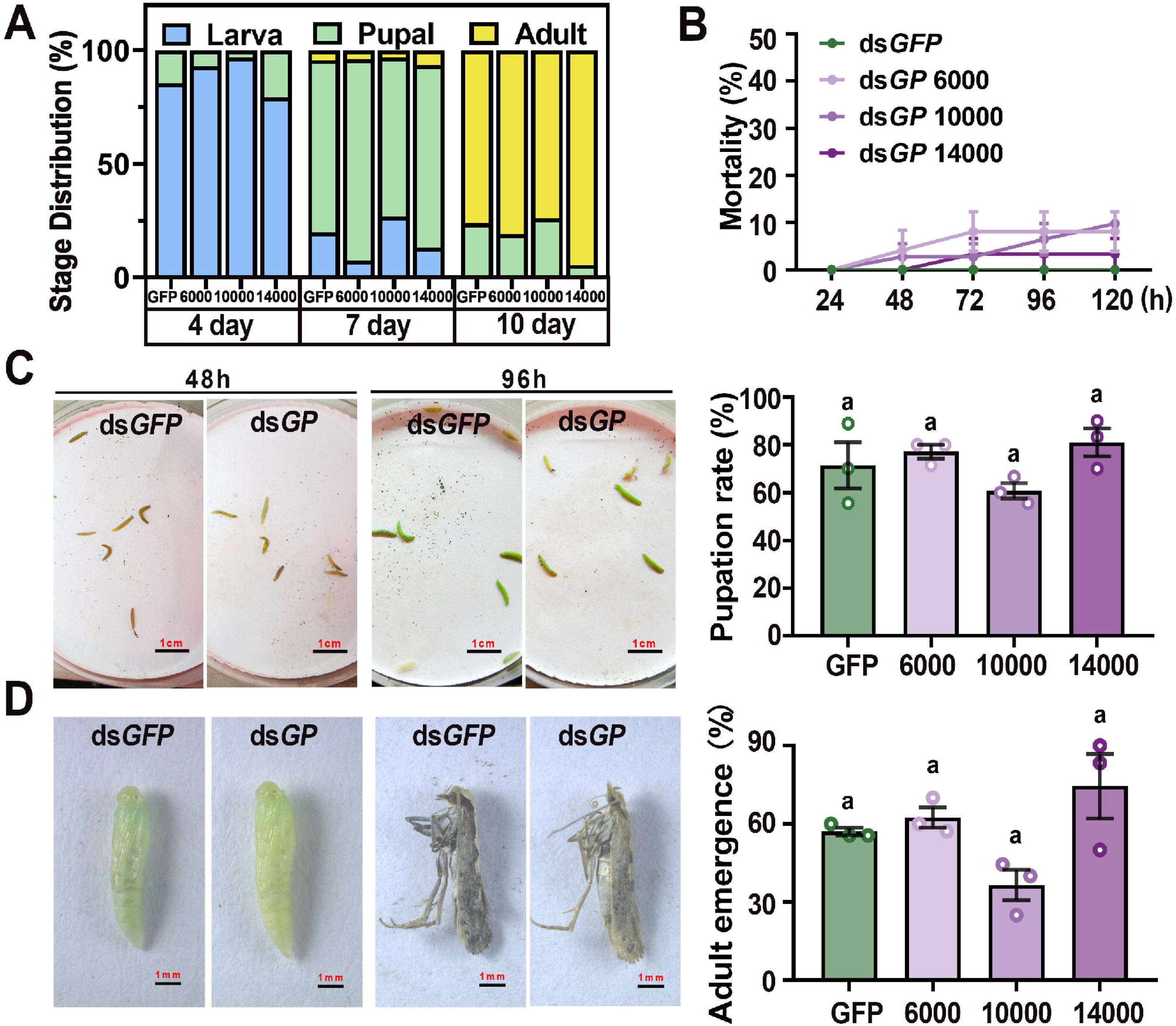
Effects of *PxGP* knockdown on GP enzyme activity and the expression of glycogen metabolism-related genes. (A) GP enzyme activity normalized to sample protein concentration. A significant decrease in activity was detected only at 48 h (*P* < 0.01). (B) GP enzyme activity per insect. Significant decreases were detected at 24 and 48 h (\**P* < 0.05, \*\**P* < 0.01). (C) Expression levels of the *GBE* gene at different time points after RNAi treatment. (D) Expression levels of the α*-Amylase* gene at different time points after RNAi treatment. All data are presented as mean ± SEM (n = 3); \**P* < 0.05, \*\**P* < 0.01, \*\*\**P* < 0.001, ds*GFP* vs. ds*GP* treatment groups. An independent samples t-test was used in (A) and (B), and two-way ANOVA with Sidak’s test was used in (C) and (D).

When GP activity was normalized to total protein content, a significant reduction was detected only at 48 h (−10.35%, *P* < 0.01; Figure 10A). However, total protein concentration itself declined substantially—by 30.78% at 24 h—before gradually recovering over subsequent time points (Figure 10–figure supplement 2). This concurrent protein decline explains the modest per-protein activity reduction: as the protein pool shrinks, the specific activity calculation is inflated even if total enzyme function is diminished. Note that the total-protein data shown in Figure 9D and in Figure 10–figure supplement 2 derive from independent experimental cohorts processed at different homogenization ratios; absolute protein concentrations are therefore not directly comparable between the two panels, and the time point at which a significant reduction was detected differs (72 h in Figure 9D; 24 h in Figure 10– figure supplement 2). Both datasets nonetheless show a transient decline of approximately 30% in total protein in ds*GP*-treated larvae within the first 72 h.

To address this, we also calculated GP activity on a per-larva basis. Per-larva GP-a activity was significantly reduced at both 24 h (−29.13%, *P* < 0.01) and 48 h (−29.28%, *P* < 0.05; Figure 10B), confirming functional suppression of GP at the protein level. Activity partially recovered at 72 and 96 h, consistent with the transcript recovery pattern observed in our previous RNAi experiments.

The 30.78% reduction in total protein at 24 h post-injection provides independent biochemical evidence supporting protein catabolism—consistent with the mobilization of amino acids to fuel gluconeogenesis as a compensatory response to GP suppression.

### *PxGP* knockdown selectively upregulates glycogen branching enzyme expression

To investigate whether alternative glycogen metabolic pathways are activated following GP suppression, we measured the expression of glycogen branching enzyme (*GBE*) and α*-amylase* by RT-qPCR in ds*GP*-treated versus ds*GFP*-treated larvae at 24, 48, 72, and 96 h post-injection (Figure 10C, D).

*GBE*, which catalyzes the formation of branch points in glycogen to increase its solubility and accessibility for degradation [37], was significantly upregulated at 24 h (+29.24%, *P* < 0.05), 48 h (+16.78%, *P* < 0.05), and 96 h (+44.46%, *P* < 0.001; Figure 10C). In contrast, α*-amylase*, a glycoside hydrolase that catalyzes hydrolysis of α-(1,4)-glycosidic bonds [38], showed no significant change at any time point (Figure 10D).

This differential expression pattern—selective *GBE* upregulation with unchanged α*-amylase* expression—indicates that the compensatory response to GP suppression involves targeted remodeling of glycogen structure rather than a generalized upregulation of all glycogen-degrading enzymes. The enhanced glycogen branching may facilitate alternative routes of glycogen mobilization even when GP-mediated phosphorolysis is impaired. However, as these observations are limited to the transcript level, further enzymatic activity assays or glycogen structure analyses would be required to determine whether this transcriptional change translates into functional alterations in glycogen mobilization.

### *PxGP* knockdown causes transient fitness costs that are fully compensated before pupation

To determine whether the metabolic changes observed following *PxGP* knockdown are associated with measurable physiological consequences, we assessed multiple fitness parameters following *PxGP* knockdown (Figure 11A–F).

**Figure 11.**
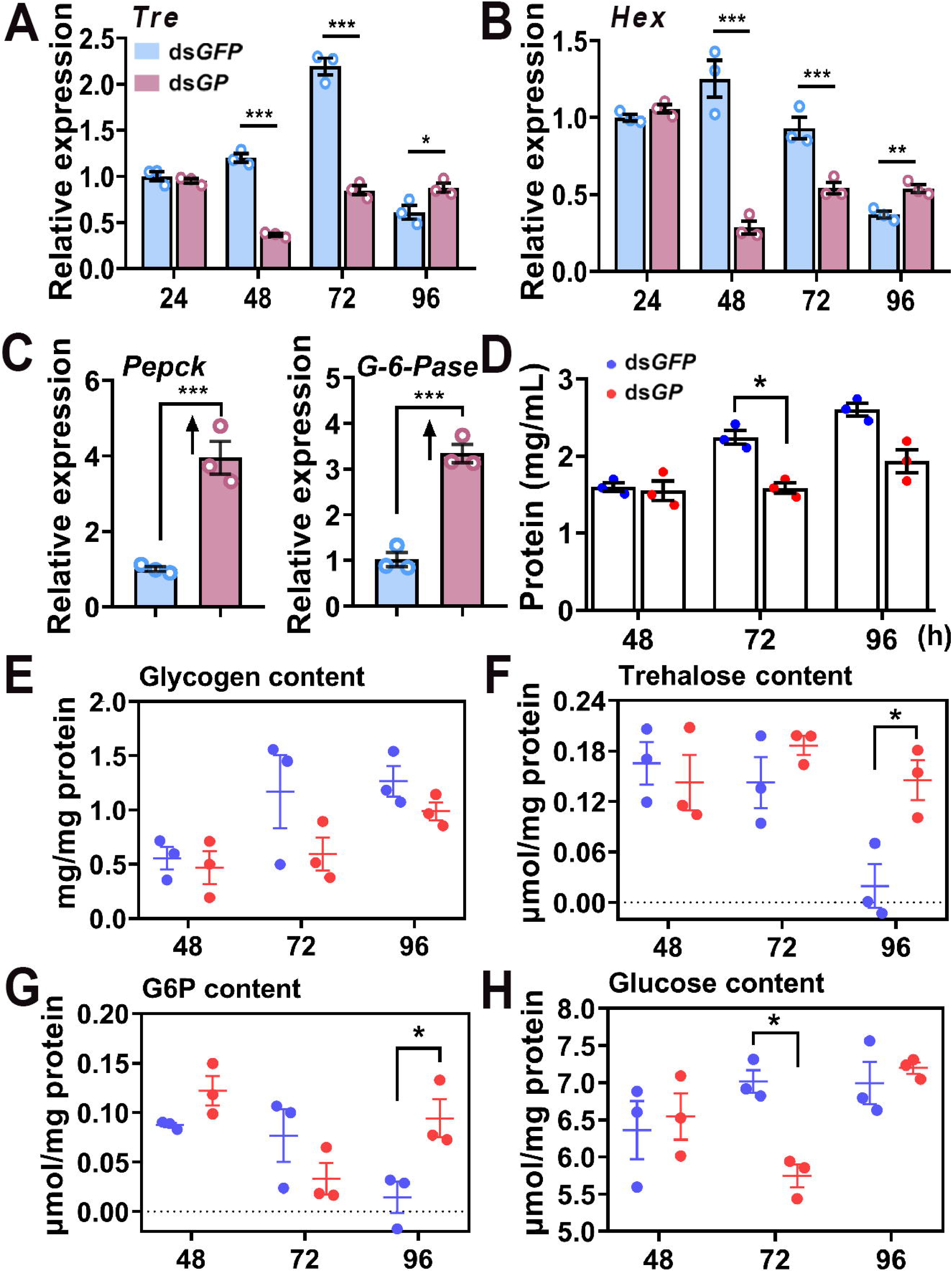
Assessment of fitness cost of *PxGP* gene silencing by RNAi in *Plutella xylostella*. (A) Schematic diagram of fitness cost evaluation at different developmental stages of *P. xylostella* after dsRNA injection. (B) Food consumption per 10 larvae at different time points. (C) Average body weight per larva at different time points. (D) Pupal weight on day 3 of pupation (n = 21). (E) Adult wing area (n = 20–21). (F) Number of eggs laid by females within 3 days (n = 10–11). Data in (B) and (C) are mean ± SEM of three biological replicates. Data in (D–F) are shown as box plots (box, interquartile range; horizontal line, median; individual points, single insects), with sample sizes as given above for each panel. \**P* < 0.05, ds*GP* vs. ds*GFP* treatment group (independent samples t-test).

Feeding rate was not significantly different between ds*GP* and ds*GFP* groups at any time point from 24 to 120 h post-injection (Figure 11B), confirming that the metabolic changes observed are independent of food intake. Larval body weight was significantly reduced in the ds*GP* group at 24 h (−29.10%, *P* < 0.05) and 48 h (−25.38%, *P* < 0.05; Figure 11C), indicating a transient but measurable physiological cost associated with metabolic compensation. However, pupal weight measured on day 3 after pupation showed no significant difference between groups (Figure 11D), demonstrating that larvae fully recover from the transient weight deficit before the pupal transition. Adult wing area (Figure 11E, Figure 11–figure supplement 1 and 2) and female 3-day fecundity (Figure 11F) were also not significantly affected.

This pattern—transient larval weight loss followed by complete recovery of pupal weight, adult morphology, and reproductive performance—is consistent with our metabolic compensation model: GP suppression likely triggers protein catabolism to supply amino acids for gluconeogenesis, contributing to transient weight loss. In the presence of continuous dietary carbohydrate supply, the compensatory response appears sufficiently effective to restore metabolic homeostasis before developmentally critical transitions.

### Molecular docking reveals the structural basis for selective inhibition of PxGP by GPI

To provide structural insight into the observed selectivity—in which GPI potently inhibits PxGP (IC_50_ = 2.96 nM) while DFB does not—we performed molecular docking and binding free energy analysis using a PxGP homology model (Figure 12, Table 1).

**Figure 12.**
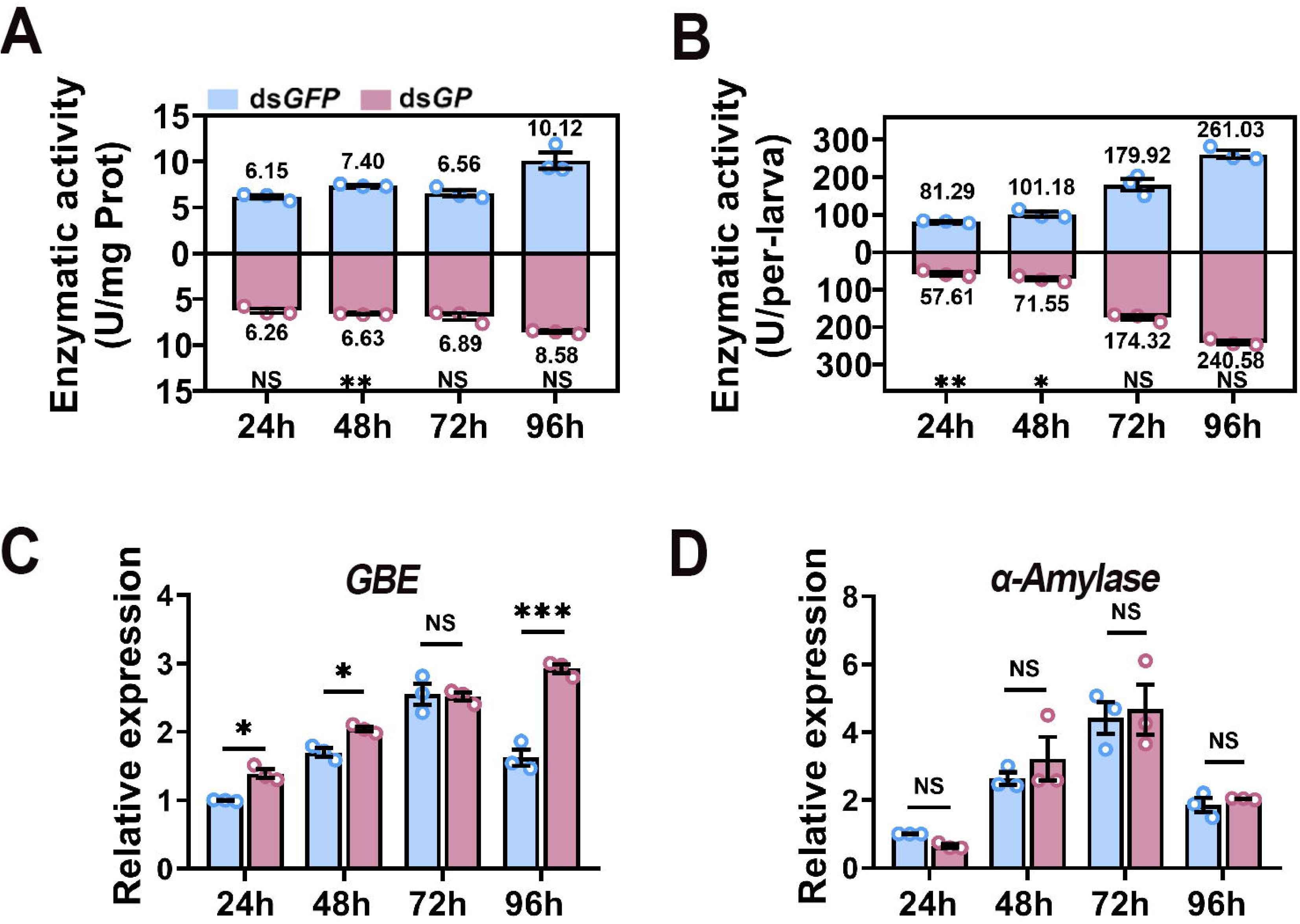
Binding mode analysis of GPI and DFB with PxGP. (A) 3D binding model of PxGP with GPI. (B, C) The key intermolecular interactions for GPI (B) and DFB (C). 2D interaction profile analysis of GPI (D) and DFB (E) with PxGP.

**Table 1.**
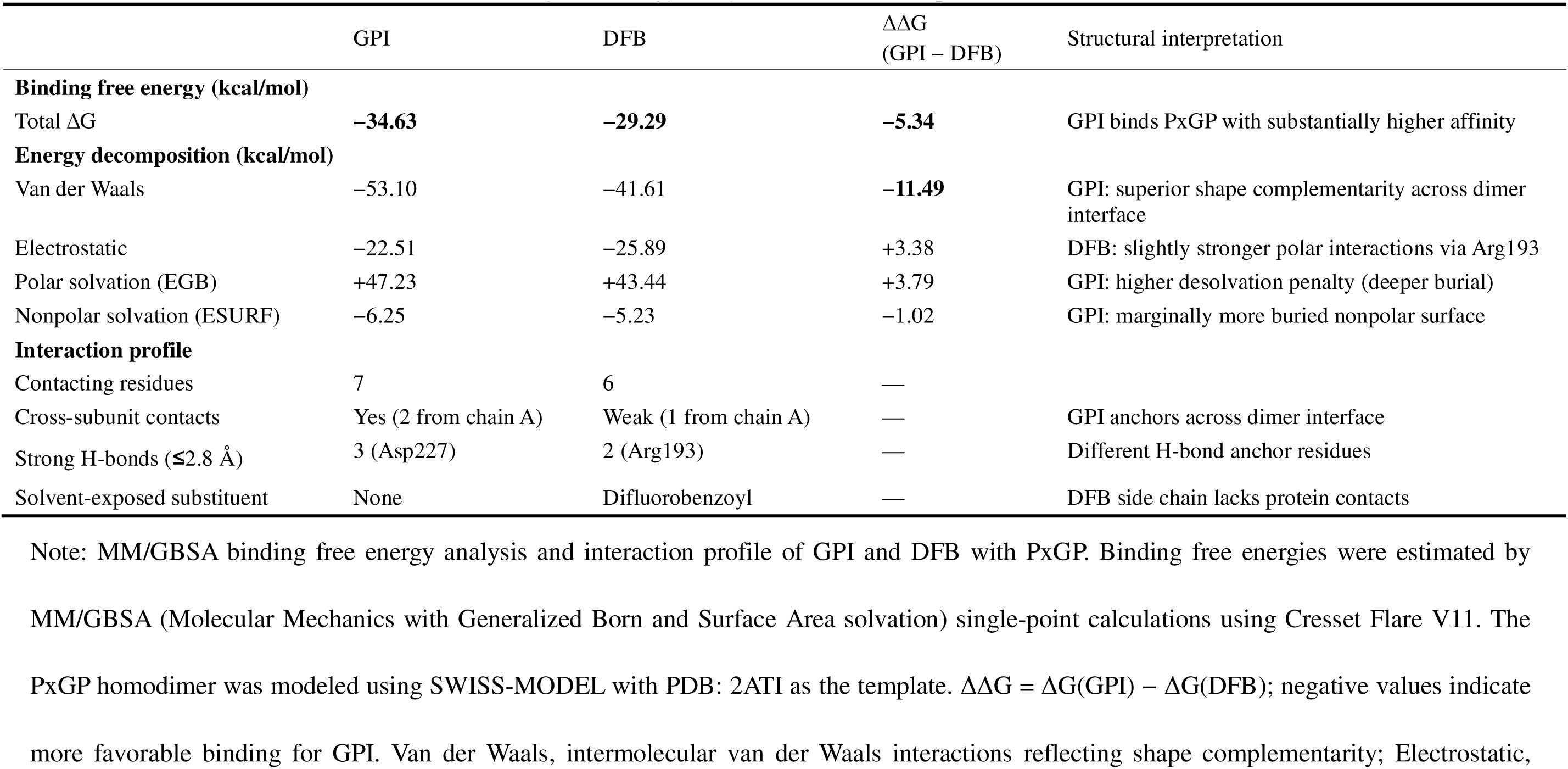
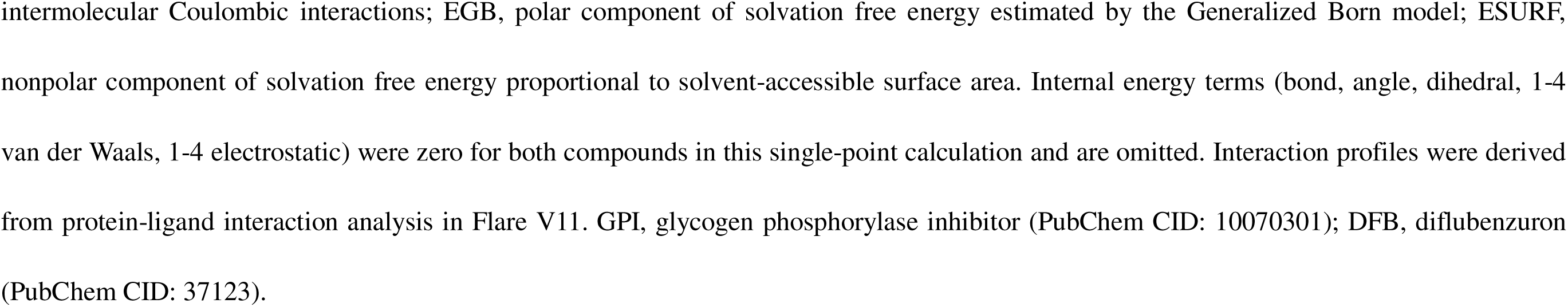
MM/GBSA binding free energy analysis and interaction profile of GPI and DFB with PxGP.

| | GPI | DFB | $\Delta\Delta G$<br>(GPI – DFB) | Structural interpretation |
| --- | --- | --- | --- | --- |
| **Binding free energy (kcal/mol)** |  |  |  |  |
| Total $\Delta G$ | **–34.63** | **–29.29** | **–5.34** | GPI binds PxGP with substantially higher affinity |
| **Energy decomposition (kcal/mol)** |  |  |  |  |
| Van der Waals | –53.10 | –41.61 | **–11.49** | GPI: superior shape complementarity across dimer interface |
| Electrostatic | –22.51 | –25.89 | +3.38 | DFB: slightly stronger polar interactions via Arg193 |
| Polar solvation (EGB) | +47.23 | +43.44 | +3.79 | GPI: higher desolvation penalty (deeper burial) |
| Nonpolar solvation (ESURF) | –6.25 | –5.23 | –1.02 | GPI: marginally more buried nonpolar surface |
| **Interaction profile** |  |  |  |  |
| Contacting residues | 7 | 6 | — |  |
| Cross-subunit contacts | Yes (2 from chain A) | Weak (1 from chain A) | — | GPI anchors across dimer interface |
| Strong H-bonds ($\leq 2.8$ Å) | 3 (Asp227) | 2 (Arg193) | — | Different H-bond anchor residues |
| Solvent-exposed substituent | None | Difluorobenzoyl | — | DFB side chain lacks protein contacts |

Note: MM/GBSA binding free energy analysis and interaction profile of GPI and DFB with PxGP. Binding free energies were estimated by
MM/GBSA (Molecular Mechanics with Generalized Born and Surface Area solvation) single-point calculations using Cresset Flare V11. The
PxGP homodimer was modeled using SWISS-MODEL with PDB: 2ATI as the template. $\Delta\Delta G = \Delta G(\text{GPI}) - \Delta G(\text{DFB})$ ; negative values indicate
more favorable binding for GPI. Van der Waals, intermolecular van der Waals interactions reflecting shape complementarity; Electrostatic,
intermolecular Coulombic interactions; EGB, polar component of solvation free energy estimated by the Generalized Born model; ESURF, nonpolar component of solvation free energy proportional to solvent-accessible surface area. Internal energy terms (bond, angle, dihedral, 1-4 van der Waals, 1-4 electrostatic) were zero for both compounds in this single-point calculation and are omitted. Interaction profiles were derived from protein-ligand interaction analysis in Flare V11. GPI, glycogen phosphorylase inhibitor (PubChem CID: 10070301); DFB, diflubenzuron (PubChem CID: 37123).

The PxGP homodimer was modeled using SWISS-MODEL with the human liver GP–acyl urea co-crystal structure (PDB: 2ATI; [15]) as the template. Molecular docking and MM/GBSA calculations were performed using Cresset Flare V11.

GPI was predicted to bind at the allosteric site located at the dimer interface, engaging seven residues across both subunits: Asn44 and Val45 from subunit A, and Trp67, Gln71, Tyr75, Arg193, and Asp227 from subunit B (Figure 12B, D). Three strong hydrogen bonds (≤ 2.8 Å) were formed with Asp227. This cross-subunit binding mode is consistent with the experimentally determined binding site of acyl urea inhibitors in the mammalian GP crystal structure [15].

DFB contacted six residues (Val45 from chain A; Ile68, Gln72, Tyr155, Arg193, and Arg242 from chain B) and formed two hydrogen bonds with Arg193 (2.1 and 2.7 Å; Figure 12C, E). Critically, DFB’s difluorobenzoyl moiety remained entirely solvent-exposed without productive protein contacts, explaining its inability to achieve effective target engagement.

MM/GBSA analysis indicated GPI’s substantially more favorable binding (ΔG = −34.63 vs. −29.29 kcal/mol; ΔΔG = −5.34 kcal/mol; Table 1). Energy decomposition revealed van der Waals interactions as the primary driver of selectivity (Δ_VDW_ = −11.49 kcal/mol), consistent with the superior shape complementarity of GPI within the dimer interface pocket.

These structural predictions are consistent with established structure–activity relationships of acyl urea compounds [15], in which target selectivity is governed by the side-chain substitution pattern rather than by the shared acyl urea core.

## Discussion

### Biochemical evidence excluding GP as a DFB target

A central challenge in mechanism-based insecticide development is distinguishing potent *in vitro* biochemical inhibitors from physiologically effective *in vivo* toxicants. This study addresses this challenge by investigating the debated molecular target of BPU insecticides. For over 40 years, the field has contended with conflicting evidence: while genetic studies implicated chitin synthase [1], direct biochemical assays consistently failed to validate it as a direct BPU target [39–41]. It has been hypothesized that BPUs might inhibit insect GP, thereby limiting glucose-derived precursors for chitin synthesis. Our systematic investigation demonstrates that while a human GPI potently inhibits PxGP, the BPU insecticide diflubenzuron does not. This discrepancy is attributable to key substitutions in its aromatic halogen substituents, which preclude effective binding to GP’s active or allosteric sites. This provides the first direct biochemical evidence excluding GP as a candidate molecular target for diflubenzuron. Consequently, BPU lethality must arise from a different mechanism, likely involving interference with chitin assembly, cuticle organization, or CHS trafficking rather than direct enzymatic inhibition, a mechanism that remains to be conclusively identified [19, 40, 41].

It is important to delimit the scope of this conclusion. Our enzymatic and structural analyses establish that diflubenzuron, specifically, does not productively engage PxGP; they do not, by themselves, exclude every member of the structurally diverse BPU class as a potential GP ligand. BPU insecticides differ substantially in their aromatic halogen substituents and, in several cases, in additional ether or alkyl side chains (Figure 1), and our MM/GBSA analysis indicates that it is precisely these side-chain architectures—rather than the shared acylurea core—that dictate whether a given acyl urea can occupy the GP allosteric pocket (Table 1). Extending the present conclusion from diflubenzuron to the BPU class as a whole would therefore require systematic testing of additional compounds. The structural framework established here (Table 1) provides a rational basis for prioritizing such an effort, by predicting which acyl urea side-chain structures are sterically and energetically compatible with productive GP binding.

### Metabolic plasticity and gluconeogenic compensation underlie GP non-essentiality

The exclusion of GP as a DFB target raised a more fundamental question: despite being a potent GP inhibitor *in vitro*, the GPI exhibited no insecticidal activity *in vivo*. This paradox—wherein a nanomolar-potency inhibitor of a putatively essential metabolic enzyme causes no phenotype—demanded mechanistic investigation.

Our investigation shows that this tolerance reflects a robust, developmentally-coordinated metabolic compensation mechanism (Figure 13). Even acute RNAi-mediated suppression of *PxGP* expression (87.59% knockdown) induced no discernible phenotype, as insects elicited a compensatory gluconeogenic pathway. This was evidenced by the concerted suppression of downstream glycogenolytic enzymes and the significant upregulation of the key gluconeogenic enzymes *PEPCK* (3.95-fold) and *G-6-Pase* (3.34-fold) at 96 h post-knockdown. Our new experimental data substantially strengthen this evidence: GP enzyme activity measurements confirm that RNAi-mediated transcript suppression translates to functional enzyme suppression *in vivo* (per-larva GP-a activity reduced by ∼29% at 24–48 h). The concurrent 30.78% decline in total protein provides independent biochemical support for protein catabolism. Furthermore, the selective upregulation of glycogen branching enzyme (GBE)—but not α-amylase—reveals a targeted compensatory response within the glycogen remodeling pathway. Fitness assessment demonstrates that this compensation, while carrying a transient larval weight cost (24–48 h), is ultimately effective: pupal weight, adult wing morphology, and female fecundity are fully maintained. This response provides a metabolic “escape route,” synthesizing glucose *de novo* from non-carbohydrate precursors to bypass the blocked glycogenolytic pathway and maintain homeostasis, particularly during the high carbohydrate demand preceding metamorphosis. Future work utilizing transcriptomic and proteomic approaches will be essential to delineate the signaling pathways (potentially involving nutrient sensors like AMPK and transcription factors such as FOXO) that link GP suppression to this precise reprogramming of metabolic gene expression [42, 43].

**Figure 13.**
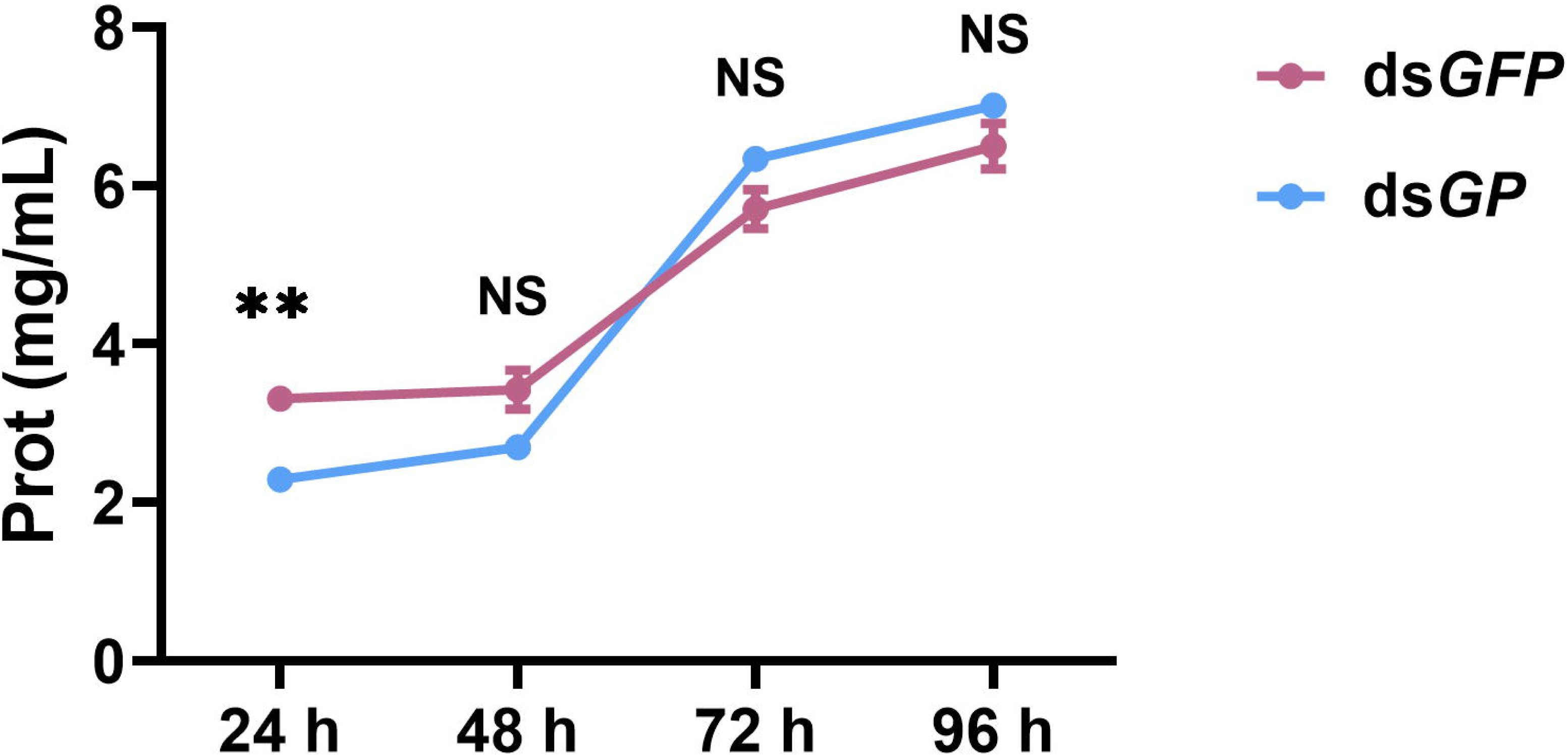
Schematic of glycogen metabolism and compensatory glucose homeostasis. Red arrows: blocked flux from glycogen to G6P; Green arrows: activated gluconeogenesis. Solid arrows represent single enzymatic steps, while dashed arrows indicate multi-step processes.

Direct metabolite quantification reflected this compensation and revealed its sophisticated temporal dynamics. The most striking finding was the lack of glycogen accumulation upon *GP* knockdown, a paradox that could be explained by the activation of GP-independent glycogen catabolism via alternative enzymes like α-amylase (which hydrolyzes internal α-1,4-bonds) and glycogen debranching enzymes (which hydrolyze α-1,6-branch points) [32–34]. However, the selective upregulation of *GBE* (rather than α*-amylase*) argues against the activation of alternative catabolic routes. Instead, reduced flux into glycogen synthesis—possibly through feedback inhibition of glycogen synthase—represents a more plausible explanation, and this will be a key focus of future investigations. At 96 h, trehalose and G6P levels in ds*GP*-treated larvae were maintained at markedly higher levels than in ds*GFP* controls, which had declined by this time point, coinciding with 3–4-fold upregulation of *PEPCK* and *G-6-Pase*. These changes strongly support, albeit indirectly, de novo glucose synthesis as the primary rescue mechanism. This biphasic response—initial metabolic stress followed by delayed (48-72 h) and robust compensation—creates a temporal buffer by 96 h and is consistent with the absence of overt phenotypes despite substantial GP suppression. This temporal coordination raises the possibility of a developmentally programmed response, potentially involving hormonal regulation, that anticipates energy demands at critical transitions. Critically, this pattern suggests a potential vulnerability window (∼72 h) during which combined inhibition of glycogenolysis and gluconeogenesis might overcome compensation, although this remains speculative and would require experimental validation.

We also acknowledge that the absence of a severe phenotype may reflect an additional, non mutually exclusive explanation: under our experimental conditions, with continuous dietary carbohydrate availability, GP activity may not be rate limiting for maintaining glucose homeostasis. The observed transient larval weight loss could thus reflect both active metabolic compensation and the inherently low demand on GP for glucose supply in a feeding larva. Distinguishing between active compensation and the non limiting nature of GP will require future studies under nutrient restricted conditions or during fasting intervals, where GP’s role is likely to become more critical.

### Implications for insecticide target selection: metabolic redundancy and the dual-target strategy

Our findings offer direct and actionable insights for the rational design of metabolism-targeting insecticides. A key lesson is that the viability of a metabolic enzyme as an insecticidal target is determined not solely by its biochemical function or pathway position but by the presence of compensatory metabolic routes. GP appears to fail as a standalone target, at least in part because of compensation via gluconeogenesis. This contrasts sharply with other carbohydrate-metabolizing enzymes that have proven to be highly effective insecticidal targets. For example, trehalase inhibitors such as Validamycin A are potent insecticides [44, 45], suggesting that trehalase inhibition may not be subject to the same degree of metabolic compensation. This distinction informs a refined strategy: dual-target inhibition. Simultaneously blocking both glycogenolysis (via GP) and gluconeogenesis (e.g., via PEPCK or G-6-Pase) would create irreversible glucose starvation, a “metabolic pincer” attack that might overcome adaptive compensation—a strategy with precedent in cancer therapy targeting both glycolysis and oxidative phosphorylation [46, 47]. Ultimately, our work highlights the value of systems-level metabolic modeling for target prioritization. Enzymes at metabolic branch points with redundant outputs are less vulnerable than those in non-bypassable pathways (e.g., chitin synthase, trehalase). Prospective application of computational flux balance analysis and constraint-based modeling, as supported by existing literature, can systematically identify such non-compensatable nodes for effective target selection [48].

### GPI concentrations and pharmacokinetic considerations

The GPI concentrations used in our larval exposure experiments (250–500 mg/L) encompass a wide dose range. Even at the highest concentration (500 mg/L), which is approximately 409,000-fold above the in vitro IC_50_ of 2.96 nM, no toxicity was observed. This apparent discrepancy between potent in vitro inhibition and absent *in vivo* toxicity could reflect several pharmacokinetic factors: limited oral bioavailability due to incomplete absorption across the midgut epithelium; metabolic inactivation by larval detoxification enzymes (cytochrome P450s, esterases, glutathione S-transferases); or sequestration by hemolymph binding proteins [49–51]. However, the metabolic phenotype we observe—elevated trehalose, reduced protein, and upregulation of gluconeogenic enzymes—provides indirect evidence that GPI does reach its target, as these changes are consistent with GP inhibition. The most parsimonious interpretation is that GPI achieves sufficient target engagement to partially suppress GP activity *in vivo*, but that the resulting metabolic compensation renders this suppression non-lethal. This interpretation is further supported by our RNAi data, in which direct genetic suppression of GP (bypassing all pharmacokinetic barriers) similarly fails to cause mortality.

### Study limitations and future directions

We acknowledge several limitations that point to productive future investigations. First, the transient nature of RNAi-mediated gene silencing in Lepidoptera precluded assessment of long-term or stage-specific consequences of GP loss; its role in adult flight or reproduction warrants investigation via CRISPR/Cas9 knockout or advanced dsRNA delivery systems [52–54]. Second, the generalizability of this gluconeogenic compensation mechanism across insect orders (e.g., Coleoptera, Diptera) remains to be tested, as metabolic regulation can vary with ecology and life history. Third, while our multi-layered evidence—from transcriptional upregulation to phenotypic rescue— provides robust qualitative support for the activation of gluconeogenesis, future studies employing ^13^C-isotope tracer analysis would be valuable to quantify the precise metabolic rates, identify primary carbon precursors, and further refine potential metabolic targets for insecticide development [55]. Extending this metabolic profiling to include key enzymes in amino acid metabolism (e.g., aspartate aminotransferase, glutamate aminotransferase) and the dynamics of lipid content would offer a more comprehensive understanding of the systemic metabolic reprogramming that accompanies GP loss.

A further consideration concerns the contribution of dietary carbohydrates to the glucose homeostasis we observed. Because our assays were performed on actively feeding larvae—and food consumption did not differ between ds*GP* and ds*GFP* groups (Figure 11B)—ingested dietary carbohydrate was continuously available and may have helped sustain hemolymph glucose alongside endogenous gluconeogenesis. Although the equivalent feeding rates argue that the metabolic phenotype is not a secondary consequence of altered food intake, our data cannot fully partition the relative contributions of *de novo* glucose synthesis and dietary carbohydrate absorption to the homeostatic buffering we documented. Starvation-challenge experiments, in which GP-suppressed larvae are deprived of dietary carbohydrate, would help isolate the endogenous gluconeogenic contribution and provide additional insight into the sufficiency of metabolic compensation under nutrient-limited conditions.

### Concluding remarks

In conclusion, our biochemical data demonstrate that DFB does not inhibit PxGP, adding enzyme-level evidence to the existing genetic framework establishing CHS involvement in BPU activity. More importantly, our findings uncover a fundamental principle of insect metabolic biology: robust compensatory pathways, specifically gluconeogenesis, can buffer against the loss of key catabolic enzymes. This metabolic plasticity renders GP non-viable as a standalone insecticidal target but illuminates a potential strategic path forward: the development of dual-target inhibitors that simultaneously block glycogenolysis and gluconeogenesis. Our structural analysis provides a molecular framework for understanding why specific acyl urea side-chain structures succeed or fail as GP inhibitors, which may inform future inhibitor design. Our findings underscore the necessity of adopting a systems-level perspective in target-based insecticide discovery. Biochemical potency *in vitro* is necessary but not sufficient for *in vivo* efficacy; the latter requires consideration of metabolic network architecture, pathway redundancy, and compensatory mechanisms. As the pest management community confronts the twin challenges of insecticide resistance and environmental sustainability, this mechanistic understanding of metabolic robustness will be essential for designing the next generation of selective, effective and durable insect control agents.

## Materials and Methods

### Insects and rearing conditions

A susceptible laboratory strain of *P. xylostella* (L.), maintained for multiple generations at the College of Plant Protection, Northwest A&F University, without prior insecticide exposure, was used for all experiments. Larvae were reared on radish (*Raphanus sativus*) seedlings grown in vermiculite. Adults were provisioned with a 10% (w/v) honey solution. All insect stages were maintained in a controlled environment at 25 ± 1°C, 60 ± 5% relative humidity (RH), and under a 16:8 h (light:dark) photoperiod.

### Cell culture and maintenance

*Spodoptera frugiperda* (Sf9) cells (Gibco, CA, USA) (Cat# 11496015) were cultured in Sf-900™ III SFM medium (Gibco) supplemented with 10% fetal bovine serum (FBS; Gibco) and 1% (v/v) Antibiotic-Antimycotic solution (Gibco, Cat# 15240062). Adherent monolayer cultures were maintained in T-flasks at 27°C. For large-scale protein expression, cells were adapted to suspension culture in baffled flasks, agitated at 120 rpm at 27°C. Cells were harvested by centrifugation (4,000 rpm, 10 min, 4°C) when the density reached 5.0 × 10^6^ cells/mL. Cell pellets were flash-frozen in liquid nitrogen and stored at −80°C until use. The supplier performs authentication and mycoplasma testing as documented in the certificate of analysis available on the manufacturer’s website. The cells were used experimentally within one year of purchase and were not re authenticated in our laboratory.

### Chemicals and reagents

Diflubenzuron (DFB, 1-(4-chlorophenyl)-3-(2,6-difluorobenzoyl) urea, ≥98% purity) was purchased from Shanghai Topscience Co., Ltd. (Shanghai, China, Cat# T0109).

The Glycogen phosphorylase-IN-1 (GPI), 1-(3-(3-(2-Chloro-4,5-difluorobenzoyl)ureido)-4-methoxyphenyl)-3-methylurea (also known as compound 42 in ref [15]), was purchased from TargetMol (Shanghai, China, Cat# T37577, ≥98% purity).

Glycogen (from bovine liver, Type III), β-glycerophosphate, HEPES, imidazole, EDTA, DTT, ATP, MgCl_2_, protease inhibitor cocktail components (AEBSF, Pepstatin A, Leupeptin), and all other biochemical reagents were purchased from Sigma-Aldrich (Shanghai, China) or Sangon Biotech (Shanghai, China) and were of molecular biology grade (≥95% purity) unless otherwise specified.

### Recombinant PxGP expression and purification

The full-length coding sequence of PxGP (2,511 bp, encoding 837 amino acids; GenBank Accession No. XM_038119125.2) was amplified from *P. xylostella* third-instar larval cDNA by PCR using gene-specific primers (Supplementary file 1), and cloned into the pFastBac1 vector (Invitrogen, Carlsbad, CA, USA) using homology-based cloning (ClonExpress II One Step Cloning Kit, Vazyme, Nanjing, China). The construct was engineered to express PxGP with an N-terminal 6×His tag (HHHHHH-) immediately following the initiating methionine, under the control of the polyhedrin (PH) promoter. The sequence fidelity of the construct was verified by Sanger sequencing (Tsingke Biotechnology, Beijing, China).

The sequence-verified pFastBac1-PxGP construct was transformed into DH10Bac™ competent cells (Invitrogen) to generate recombinant bacmid DNA via Tn7-mediated transposition, following the manufacturer’s Bac-to-Bac® protocol. Recombinant bacmid DNA was isolated, verified by PCR using M13 forward and reverse primers (Supplementary file 1), and transfected into Sf9 cells using Cellfectin® II Reagent (Invitrogen) to generate P0 viral stock. The viral titer was amplified through two additional passages (P1, P2) to obtain high-titer P3 viral stock.

For large-scale protein production, Sf9 cells in suspension culture (2.0 × 10^6^ cells/mL, 500 mL culture volume in a 2-L baffled Erlenmeyer flask) were infected with P3 viral stock at a Multiplicity of Infection (MOI) of 5. Infected cultures were maintained at 27°C with agitation at 120 rpm. Cells were harvested 72 h post-infection by centrifugation (4,000 rpm, 10 min, 4°C), yielding approximately 8.25 g of cell pellet. Cell pellets were flash-frozen in liquid nitrogen and stored at −80°C until use.

For protein purification, frozen cell pellets were thawed on ice and resuspended in ice-cold Lysis Buffer (50 mM β-glycerophosphate [pH 7.5], 150 mM NaCl, 0.2 mM DTT, 0.5 mM EDTA, and 1× protease inhibitor cocktail [1 mM AEBSF, 15 µM Pepstatin A, 20 µM Leupeptin]) at a ratio of 4 mL buffer per gram of cell pellet. Cells were lysed by sonication on ice using a Scientz-IID sonicator (Ningbo Scientz Biotechnology, China) with the following parameters: 6 cycles of 30 s ON / 60 s OFF at 40% amplitude. The lysate was clarified by centrifugation (16,000 × g, 30 min, 4°C).

The clarified supernatant was loaded onto a 5-mL HisTrap™ HP column (GE Healthcare, Uppsala, Sweden) pre-equilibrated with Lysis Buffer containing 10 mM imidazole. The column was first washed with 10 column volumes (CV) of Equilibration Buffer (Lysis Buffer + 10 mM imidazole), followed by 5 CV of Wash Buffer (Lysis Buffer + 30 mM imidazole). Bound 6×His-tagged PxGP protein was eluted with a 15 CV linear gradient from 30 to 200 mM imidazole in Lysis Buffer. Fractions (2 mL each) were collected and analyzed by 13% SDS-PAGE with Coomassie Brilliant Blue R-250 staining. Fractions containing PxGP, identified as a prominent band at ∼100 kDa, were pooled and dialyzed overnight at 4°C against Storage Buffer (50 mM HEPES [pH 7.4], 150 mM NaCl, 1 mM DTT, 10% [v/v] glycerol) using a 10 kDa molecular weight cut-off dialysis membrane (Spectrum Labs, Rancho Dominguez, CA, USA), with three buffer changes. Protein concentration was determined using the Pierce™ BCA Protein Assay Kit (Thermo Fisher Scientific, Waltham, MA, USA) with bovine serum albumin (BSA) as the standard. Aliquots of purified PxGP were flash-frozen in liquid nitrogen and stored at −80°C. The typical yield was 7.5 mg of purified protein from 8.25 g of cell pellet.

### *In vitro* activation of recombinant PxGP

Purified recombinant PxGP exists predominantly in the inactive dephosphorylated b form (PxGP-b) and requires phosphorylation at a conserved serine residue (corresponding to Ser14 in mammalian GP) for conversion to the active a form (PxGP-a) [17]. Activation was performed *in vitro* using immobilized Phosphorylase Kinase (PhK) (Cat# HY-P2757, MedChemExpress, Monmouth Junction, NJ, USA).

Immobilized PhK beads (100 µL slurry) were washed three times with 1 mL of Kinase Buffer (25 mM β-glycerophosphate, pH 7.5, 0.5 mM EDTA, 0.25 mM DTT, 1× protease inhibitor cocktail) and resuspended in 500 µL of dialyzed PxGP-b solution (final protein concentration: 1 mg/mL). The activation reaction was initiated by adding ATP (sodium salt) to a final concentration of 7 mM and MgCl_2_ to 22 mM. The suspension was incubated for 2 h at room temperature (22-25°C) with gentle end-over-end rotation.

Following incubation, the activated PxGP-a (present in the supernatant) was separated from the immobilized PhK beads using a magnetic separation stand. The supernatant was passed through a 0.22 µm syringe filter (Millipore) to remove any residual beads and was used immediately for enzymatic assays. The extent of activation was assessed by measuring specific activity (see glycogen phosphorylase activity assay). Activated PxGP-a was used fresh and not stored, as phosphorylated GP is subject to dephosphorylation by endogenous phosphatases over time.

### Western blot analysis

Protein samples were separated by 13% SDS-PAGE (120 V, 90 min) and electrophoretically transferred to a nitrocellulose (NC) membrane (0.45 µm pore size, Millipore, Burlington, MA, USA) at 300 mA for 90 min at 4°C using a Bio-Rad Mini Trans-Blot® system. The membrane was blocked for 1 h at room temperature in 5% (w/v) non-fat dry milk in TBST (10 mM Tris-HCl, pH 7.4, 150 mM NaCl, 0.1% [v/v] Tween-20).

The membrane was incubated overnight at 4°C with an anti-His mouse monoclonal antibody (1:1500 dilution in blocking buffer; Cat# ABT2050, Abbkine, Wuhan, China). After three washes with TBST (10 min each), the membrane was incubated for 1 h at room temperature with an alkaline phosphatase (AP)-conjugated goat anti-mouse IgG secondary antibody (1:4000 dilution in blocking buffer; Cat# A0208, Beyotime Biotechnology, Shanghai, China). Following three additional TBST washes, immunoreactive bands were visualized using BM Purple AP Substrate (Roche, Basel, Switzerland) according to the manufacturer’s instructions. Images were captured using a ChemiDoc™ XRS+ imaging system (Bio-Rad, Hercules, CA, USA).

### Glycogen phosphorylase activity assay

GP enzymatic activity was measured using a coupled-enzyme spectrophotometric assay kit (Cat# BC3345, Solarbio Life Science, Beijing, China). This assay quantifies GP activity by measuring the GP-catalyzed phosphorolysis of glycogen to glucose-1-phosphate (G-1-P). The product G-1-P is enzymatically converted to glucose-6-phosphate (G6P) by phosphoglucomutase (PGM), and G6P is subsequently oxidized by glucose-6-phosphate dehydrogenase (G6PDH), which stoichiometrically reduces NADP^+^ to NADPH. The rate of NADPH formation, which is directly proportional to GP activity, was monitored by measuring the increase in absorbance at 340 nm (A_340_) using a Infinite 200 PRO microplate reader (Tecan, Männedorf, Switzerland).

The reaction mixture (200 µL total volume per well in a 96-well UV-transparent microplate) contained the following components (as supplied in the kit): 160 µL of Reaction Buffer (containing glycogen [2 mg/mL final], inorganic phosphate [10 mM], AMP [1 mM], NADP^+^ [0.4 mM], PGM, G6PDH, MgCl_2_, and appropriate buffer), 30 µL of deionized water, and 10 µL of enzyme sample (purified PxGP-a or crude larval lysate). The reaction was initiated by adding the enzyme sample, and A_340_ was recorded continuously at 15 s intervals for 30 min at 25°C.

Specific activity was calculated using the following formula:

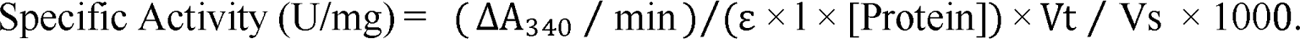

where: ΔA_340_ represents the change in absorbance at 340 nm, ε = molar extinction coefficient of NADPH at 340 nm (6.22 mM^-1^ · cm^-1^), l = light path length (0.6 cm for 200 µL in a 96-well microplate), [Protein] = protein concentration in the assay (mg/mL), Vt = total reaction volume, Vs = sample volume added.

One Unit (U) of GP activity is defined as the amount of enzyme that catalyzes the formation of 1 µmol of NADPH per minute under the assay conditions.

### Inhibitor screening and IC_50_ determination

For *in vitro* enzyme inhibition assays, purified activated PxGP-a was diluted to 10 µg/mL in Assay Dilution Buffer (30 mM HEPES, pH 7.2, 60 mM KCl, 1.5 mM EDTA, 1.5 mM MgCl_2_). Test compounds (GPI, DFB) were prepared as 100× stock solutions in DMSO and serially diluted in Assay Dilution Buffer to achieve final assay concentrations ranging from 0.16 nM to 500 nM (for GPI) and 0.54 nM to 1700 nM (for DFB), respectively (final DMSO concentration ≤ 1% [v/v] in all wells, including vehicle controls).

The enzyme (10 µL of 10 µg/mL PxGP-a, final concentration 10 µg/mL in the assay) was pre-incubated with 10 µL of inhibitor solution (or vehicle control) for 15 min at room temperature in a 96-well microplate. The reaction was then initiated by adding 180 µL of the Reaction Buffer (as described in glycogen phosphorylase activity assay), and A_340_ was monitored continuously for 30 min at 25°C.

Residual enzyme activity at each inhibitor concentration was calculated as a percentage of the vehicle control (1% DMSO alone). Dose-response curves were generated by plotting percent residual activity versus log[inhibitor]. IC_50_ values (the concentration of inhibitor required to reduce enzyme activity by 50%), Hill slopes, and 95% confidence intervals (CI) were calculated by non-linear regression analysis using a four-parameter logistic equation in GraphPad Prism 9.0 (GraphPad Software Inc., San Diego, CA, USA):

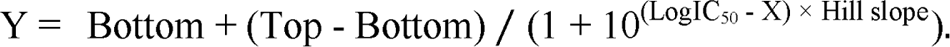

where Y is percent residual activity, X is log[inhibitor], Top and Bottom are the upper and lower plateaus of the curve (constrained to 100% and 0%, respectively), and Hill slope describes the steepness of the curve.

For each compound, at least three independent experiments were performed, each with triplicate technical replicates.

### Preparation of larval crude lysate for *in vivo* activity

To assess the inhibitory effects of GPI and DFB on native GP activity *in vivo*, crude enzyme extracts were prepared from *P. xylostella* larvae. Healthy third-instar (L3) larvae (n = 30 per biological replicate, three replicates total) were collected, briefly washed in ice-cold phosphate-buffered saline (PBS, pH 7.4) to remove surface contaminants, and gently blotted dry on filter paper. Larvae were then homogenized on ice in 3 volumes (w/v) of ice-cold Homogenization Buffer (30 mM HEPES, pH 7.2, 60 mM KCl, 1.5 mM EDTA, 1.5 mM MgCl_2_, 1× protease inhibitor cocktail).

The homogenate was centrifuged at 15,000 × g for 20 min at 4°C, and the supernatant (crude enzyme extract) was carefully collected, avoiding the lipid layer and pellet. Protein concentration was determined using the BCA assay, and extracts were either used immediately for activity assays or aliquoted and stored at −80°C (activity remained >90% for up to 2 weeks).

GP activity in crude lysates was measured using the same coupled-enzyme assay described in glycogen phosphorylase activity assay, with the following modifications: crude lysate (diluted to 2 mg/mL total protein) was used in place of purified enzyme, and pre-incubation with inhibitors was performed for 5 to 45 min to assess time-dependent inhibition. Final inhibitor concentrations tested were 10^-5^, 10^-6^ or 10^-7^ M.

### Sequence alignment and bioinformatic analysis

The amino acid sequence of PxGP (XP_037975053.2) was obtained from the NCBI database. Multiple sequence alignment with orthologs from *Homo sapiens* (NP_002854.3), *Oryctolagus cuniculus* (NP_001075653), *Helicoverpa armigera* (XP_021190210.1), *Spodoptera exigua* (ACN78408.1), and *Ostrinia furnacalis* (AFO54708.2) was performed using ClustalX. Conserved domains and motifs were analyzed using the ScanProsite and NCBI CD-Search web servers. Protein pairwise similarity matrix was visualized as a heatmap using the “HeatMap” module in TBtools-II (v2.390) [56].

### Larval bioassays

Larval bioassays were conducted using a standard leaf-dip method adapted from established protocols [57]. Stock solutions of GPI and DFB were prepared in HPLC-grade acetone at 10,000 mg/L. Serial dilutions were prepared in deionized water containing 0.1% (v/v) Triton X-100 as a wetting agent to achieve final test concentrations of 250 and 500 mg/L for GPI, and 125 and 250 mg/L for DFB. The final acetone concentration in all treatment solutions was 1% (v/v). A solvent control group (CK) consisted of 1% acetone and 0.1% Triton X-100 in water without any test compound.

Fresh cabbage (*Brassica oleracea* var. *capitata*) leaf discs (5 cm diameter) were cut using a cork borer, dipped in the respective treatment solutions for 6-8 s with gentle agitation, and then air-dried for 30 min at room temperature in a fume hood. Once dry, each leaf disc was placed in a sterile 90 mm petri dish lined with moistened filter paper.

Ten synchronized third-instar larvae of *P. xylostella* (24-36 h post-molt to L3) were transferred to each treated leaf disc using a soft brush. Each treatment consisted of three biological replicates (i.e., three Petri dishes, 30 larvae total per treatment). Petri dishes were sealed with Parafilm® and maintained under standard rearing conditions (25 ± 1°C, 60 ± 5% RH, 16:8 h L:D photoperiod). Leaf discs were replaced with freshly treated discs every 12 h to ensure continuous exposure.

Mortality was recorded every 24 h for 120 h (5 days). Larvae that failed to respond to gentle prodding or showed moribund behavior (inability to right themselves or move coordinately) were considered dead. Corrected mortality was calculated using Abbott’s formula [58]:

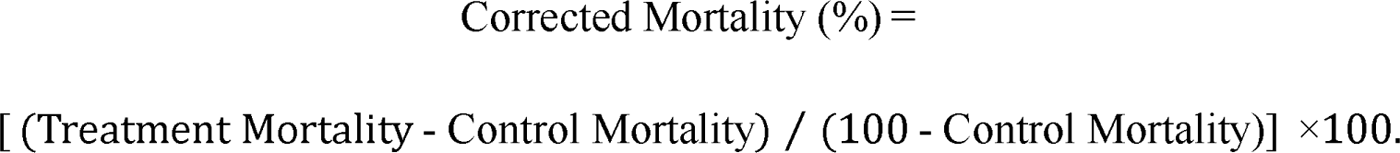

Pupation rates (number of pupae formed / initial number of larvae × 100) and adult emergence rates (number of emerged adults / number of pupae formed × 100) were recorded at 8 and 10 days post-treatment, respectively.

Lethal concentration values (LC_30_ and LC_50_) and 95% confidence intervals for DFB were calculated using probit analysis in POLO-Plus software (LeOra Software, Berkeley, CA, USA).

### GPI exposure for gene expression analysis

For time-course gene expression analysis, synchronized third-instar larvae were exposed to 500 mg/L GPI (the highest concentration tested in bioassays) or solvent control (1% acetone + 0.1% Triton X-100) using the leaf-dip method described in larval bioassays.

Surviving larvae were collected at 24, 48, 72, and 96 h post-treatment. At each time point, three biological replicates (10 pooled larvae each) were flash-frozen in liquid nitrogen and stored at −80°C until RNA extraction. This sampling scheme ensured sufficient RNA yield while minimizing inter-individual variability.

### Quantitative real-time PCR (RT-qPCR) analysis

Total RNA was extracted from frozen samples using RNAiso Plus reagent (TaKaRa, Kusatsu, Japan). RNA concentration and purity were assessed by NanoDrop 2000c spectrophotometer (Thermo Fisher Scientific). Only samples with A_260_/A_280_ ratios between 1.9 and 2.1 and A_260_/A_230_ ratios >1.8 were used for downstream analysis. RNA integrity was confirmed by agarose gel electrophoresis (sharp 28S/18S rRNA bands, ratio ≈ 2:1). cDNA was synthesized from 1 µg RNA using PrimeScript™ RT Reagent Kit with gDNA Eraser (TaKaRa).

Gene-specific primers for *PxGP*, *trehalase* (*PxTre*), *hexokinase* (*PxHex*), phosphoenolpyruvate carboxykinase (*PxPEPCK*), *PxG-6-Pase*, and the reference gene *PxRPS13* (ribosomal protein S13) were designed using Primer-BLAST (NCBI) and are listed in Supplementary file 1. *PxRPS13* was selected based on its stable expression across developmental stages and treatment conditions (geNorm M value < 0.5) [59]. All primers were validated for specificity (single peak in melt-curve), efficiency (90-110%, R² > 0.98), and absence of primer-dimers (no-template controls). RT-qPCR reactions were performed on a QuantStudio™ 5 Real-Time PCR System (Thermo Fisher Scientific) using TB Green® Premix Ex Taq™ II (Tli RNaseH Plus) (2×) (TaKaRa) in 20 µL reactions. The thermal cycling conditions were as follows: initial denaturation at 95°C for 30 s; 40 cycles of 95°C for 15 s (denaturation) and 60°C for 30 s (annealing/extension); followed by melt-curve analysis. Each sample was analyzed in technical triplicate, and each biological replicate consisted of three independent samples. Relative gene expression was calculated using the 2^-ΔΔCt^ method [60].

### RNA interference (RNAi)

A conserved 606 bp region of *PxGP* (nucleotides 66-671, part of the AMP-binding domain) was selected for RNAi. The fragment was amplified from larval cDNA using primers flanked by T7 promoters (Supplementary file 1). Double-stranded RNA (dsRNA) was synthesized with the T7 RiboMax™ Express RNAi System (Promega, Madison, WI, USA). The dsRNA of green fluorescent protein (ds*GFP*, 541 bp) was synthesized using a pGFP plasmid template as a non-specific negative control. Following synthesis, dsRNA was treated with RNase-free DNase I (30 min at 37°C), purified by isopropanol precipitation, and integrity verified by 1.5% agarose gel electrophoresis. Concentration and purity (A_260_/A_280_=1.9-2.1) was measured using a NanoDrop spectrophotometer. Purified dsRNA was resuspended in nuclease-free water at concentrations of 6,000, 10,000, or 14,000 ng/µL and stored in single-use aliquots at −80°C.

For microinjection, synchronized early third-instar larvae (6-12 h post-molt) were briefly chilled on ice for 2-3 min and injected laterally using a DMP-300 Injector (Micrology Precision Instruments, Ltd, Wuhan, China). After recovery on fresh cabbage leaves for 30 min at room temperature, larvae were maintained under standard rearing conditions. Phenotypic parameters (mortality, molting success, pupation rate, adult emergence) were monitored daily for 10 days. Larvae were collected at 24, 48, 72, and 96 h post-injection for RT-qPCR analysis to confirm target gene knockdown (n = 3 biological replicates, each containing 10 pooled larvae per time point).

### Metabolite quantification

To directly assess metabolic flux alterations following Px*GP* knockdown, we quantified four key metabolites in the glycogen-glucose metabolic axis using commercial enzymatic assay kits: glycogen and glucose (Glycogen Assay Kit, Grace Biotechnology, Suzhou, China, Cat# G0590W96), trehalose (Trehalose Assay Kit, Grace Biotechnology, Cat# G0553W96), and glucose-6-phosphate (G6P Assay Kit, Beyotime Biotechnology, Shanghai, China, Cat# S0185). All kits utilize colorimetric or fluorometric enzymatic reactions and were performed according to the manufacturers’ protocols with minor modifications as described below.

Third-instar larvae injected with ds*GP* (14,000 ng/µL, the highest dose achieving maximal knockdown) or ds*GFP* control were collected at 48, 72, and 96 h post-injection. At each time point, three biological replicates were collected, each consisting of 10 pooled larvae. Samples were immediately flash-frozen in liquid nitrogen and stored at −80°C until analysis.

For metabolite extraction, frozen larvae were homogenized on ice in ice-cold extraction buffer specific to each metabolite assay (typically 100 µL buffer per 10 mg tissue). For glycogen extraction, samples were homogenized in deionized water and immediately boiled for 3 min to inactivate endogenous glycogenolytic enzymes. For trehalose, glucose, and G6P extraction, samples were homogenized in the respective assay buffers provided in the kits. Homogenates were centrifuged at 12,000 × g for 10 min at 4°C, and supernatants were immediately used for metabolite quantification or stored at −80°C for no more than 24 h.

Metabolite quantification was performed in 96-well microplates using Infinite 200 PRO microplate reader (Tecan, Männedorf, Switzerland) at the wavelengths specified in each kit’s protocol. Sample metabolite concentrations were calculated by comparison with the absorbance of the kit’s standard. Metabolite concentrations were normalized to tissue protein content to account for variation in tissue input. Protein concentration in the crude lysates was determined using the Protein Quantification Kit (BCA Assay) (Abbkine Scientific) with bovine serum albumin as the standard. All samples were analyzed in technical triplicate. Final metabolite levels were expressed as: Glycogen: mg/mg protein; Glucose: µmol/mg protein; Trehalose: µmol/mg protein; G6P: µmol/mg protein. Because metabolite levels are expressed per unit total protein, the changes in total protein content that accompany *PxGP* knockdown bear directly on their interpretation. Total protein declined substantially at the earlier time points (up to ∼30%; Figure 9D, Figure 10–figure supplement 2) before recovering, which means that for any metabolite whose absolute amount was unchanged, the protein-normalized value would be correspondingly inflated during the protein-depleted window. The protein-normalized metabolite data reported here should therefore be interpreted with this denominator effect in mind. We retained protein normalization because it remains the most appropriate correction for the variation in tissue input inherent to pooled-larvae sampling; we note, however, that absolute (per-larva) quantification — as we additionally applied to GP enzyme activity (Figure 10A, B) — provides a complementary, denominator-independent perspective and represents a useful refinement for future metabolite profiling.

### Glycogen phosphorylase activity assay in RNAi-treated larvae

To assess GP enzyme activity following RNAi, early third-instar larvae were injected with 14,000 ng/μL ds*GP* or ds*GFP*. At 24, 48, 72, and 96 h post-injection, 10 larvae per replicate (three biological replicates per time point) were collected. Active glycogen phosphorylase a (GP-a) activity was measured using a coupled-enzyme spectrophotometric assay kit (catalog no. BC3345, Solarbio, Beijing, China) following the manufacturer’s protocol as described above. GP activity was normalized both to total protein content (U/mg protein) and to individual larva (U/larva).

### RT-qPCR analysis of alternative glycogen metabolic enzymes

Homologs of human glycogen branching enzyme (GBE, CAG9111719.1), *Pieris rapae* α-amylase (XP_011548197.3), and glycogen debranching enzyme (GDE, XP_048484812.1) were identified in the *P. xylostella* genome database by sequence homology search. RT-qPCR primers were designed using Primer-BLAST (NCBI) (Supplementary file 1). Multiple primer pairs designed for *GDE* failed to yield amplification products; therefore, only *GBE* and α*-amylase* were analyzed. *PxRPS13* (ribosomal protein S13) was used as the reference gene. Total RNA was extracted from larvae collected at 24, 48, 72, and 96 h post-injection (three biological replicates, 10 larvae per replicate). Relative expression levels were calculated using the 2^−ΔΔCt^ method and analyzed by two-way ANOVA with Šidák’s multiple comparisons test [61].

### Fitness cost assessment

To evaluate fitness consequences of *PxGP* knockdown, the following parameters were measured. Feeding rate: cabbage leaf discs of uniform size were provided every 12 h, and remaining leaf area was measured by ImageJ at 24 h intervals from 24 to 120 h post-injection. Larval weight: 10 larvae per replicate were weighed at 24 h intervals. Pupal weight: all pupae were weighed on day 3 after pupation. Adult wing area: forewings were photographed using a SteREO Discovery V20 stereomicroscope (Zeiss, Germany) and measured using ImageJ. Female fecundity: newly emerged females were paired with untreated males, and eggs deposited on leaf discs were counted for 3 consecutive days. Three biological replicates were performed. Data were analyzed by independent-samples t-test.

### Homology modeling and molecular docking

The three-dimensional structure of PxGP was generated by homology modeling using SWISS-MODEL (https://swissmodel.expasy.org [62]) with the crystal structure of human liver glycogen phosphorylase a in complex with an acyl urea inhibitor (PDB: 2ATI [15]) as the template. Since PxGP functions as a homodimer, the dimeric biological assembly was constructed based on the template’s quaternary structure. The three-dimensional structures of GPI (PubChem CID: 10070301) and DFB (PubChem CID: 37123) were obtained from PubChem in SDF format. Molecular docking was performed using Cresset Flare V11 (Cresset, Litlington, UK). Binding free energies were estimated by MM/GBSA (Molecular Mechanics with Generalized Born and Surface Area solvation) single-point calculations. Two-dimensional protein-ligand interaction diagrams were generated within Flare V11.

### Use of artificial intelligence tools

AI-based language editing tools were used to assist with English grammar and style refinement during manuscript preparation. All scientific content, including experimental design, data collection, analysis, and interpretation, was generated entirely by the authors.

## Supporting information

Primer sequences used in this study.

Toxicity of Diflubenzuron (DFB) against 3rd-instar Plutella xylostella larvae at 120 h.

---

**Figure 2–figure supplement 1.**
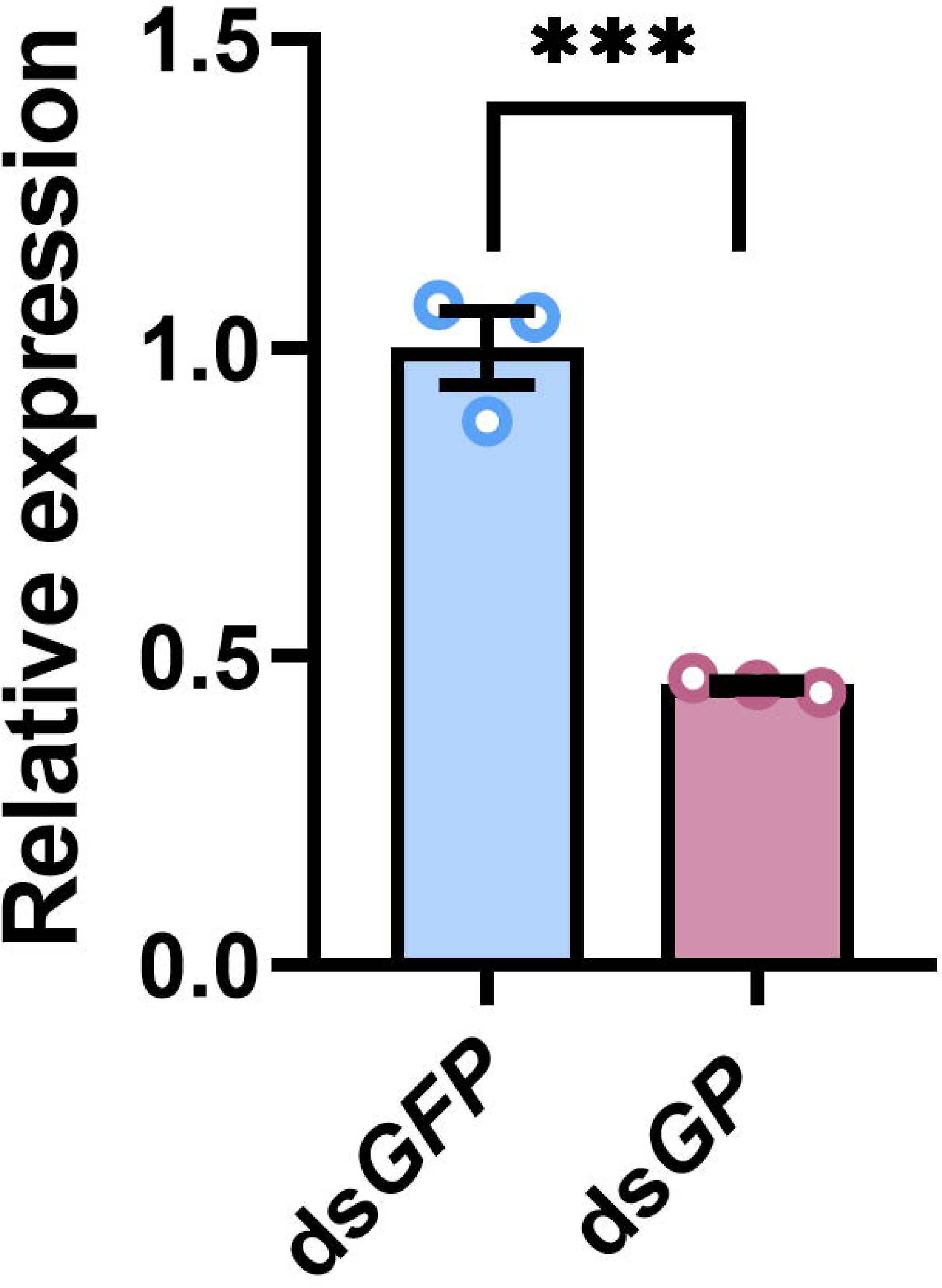
Enzymatic activity of purified PxGP and inhibition assays using crude larval lysate. (A) Reaction kinetics of purified, activated PxGP-a (red line) compared to a no-enzyme control (CK, blue line). The specific activity was calculated to be 294.25 ± 3.45 U/mg Prot (mean ± SEM). (B) Inhibition of native GP activity in a crude larval lysate by GPI. Activity was measured at 5, 15, 25, and 45 min post-incubation. (C) Effect of DFB on native GP activity in a crude larval lysate. DFB showed no significant inhibition at any concentration or time point. Data in (B) and (C) are presented as mean ± SEM (n = 3).

**Figure 5–figure supplement 1.**
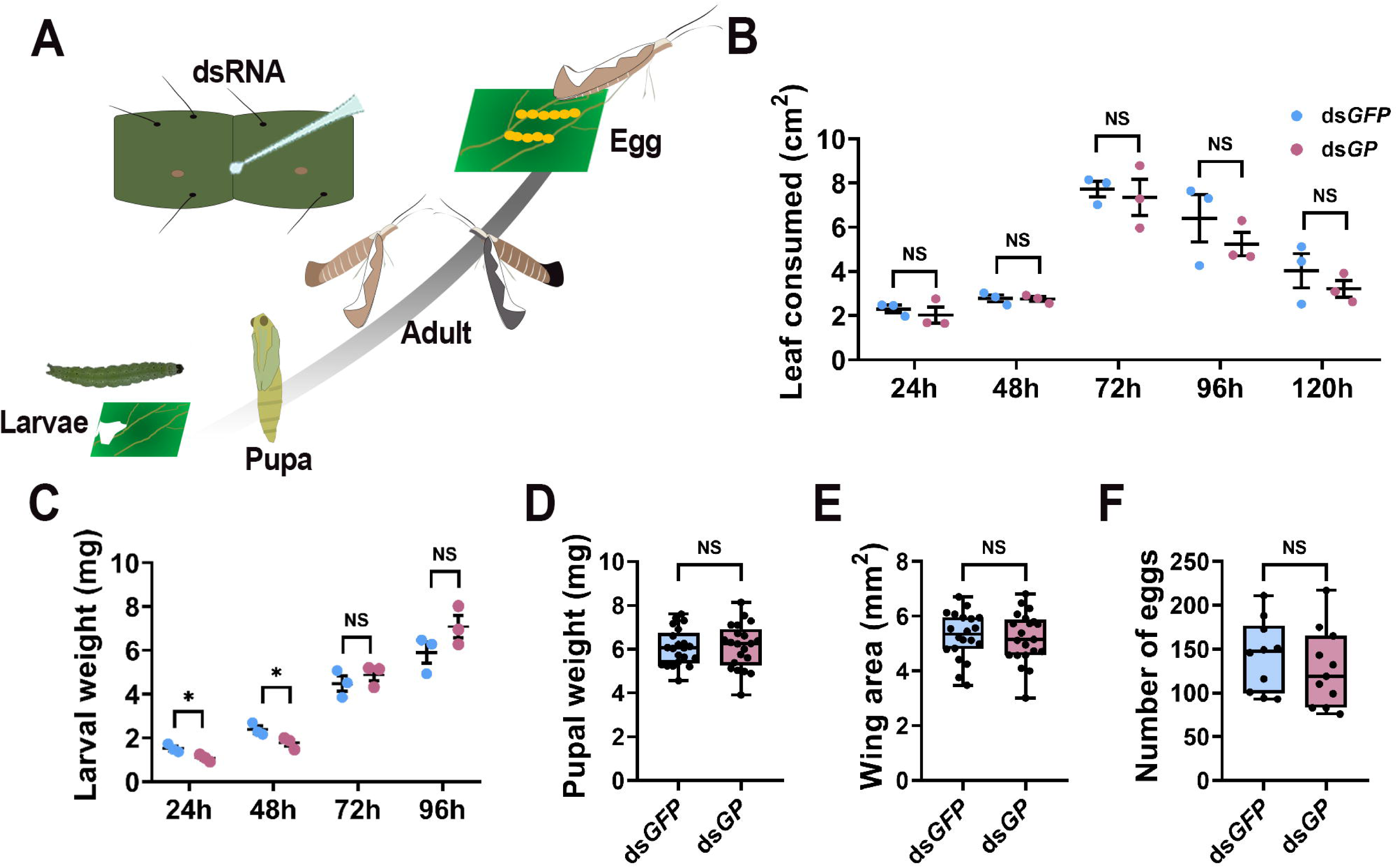
Heatmap depicting pairwise sequence similarity of glycogen phosphorylase among six species. The heatmap displays the percentage of global sequence similarity calculated from pairwise alignments using TBtools. The six species (*Plutella xylostella*, *Homo sapiens*, *Oryctolagus cuniculus*, *Helicoverpa armigera*, *Ostrinia furnacalis* and *Spodoptera exigua*) are arranged in the same order along both axes. The color intensity in each cell corresponds to the similarity percentage, as shown in the color key (from blue (low) to red (high)). Numerical values are shown within the cells.

**Figure 10–figure supplement 1.**
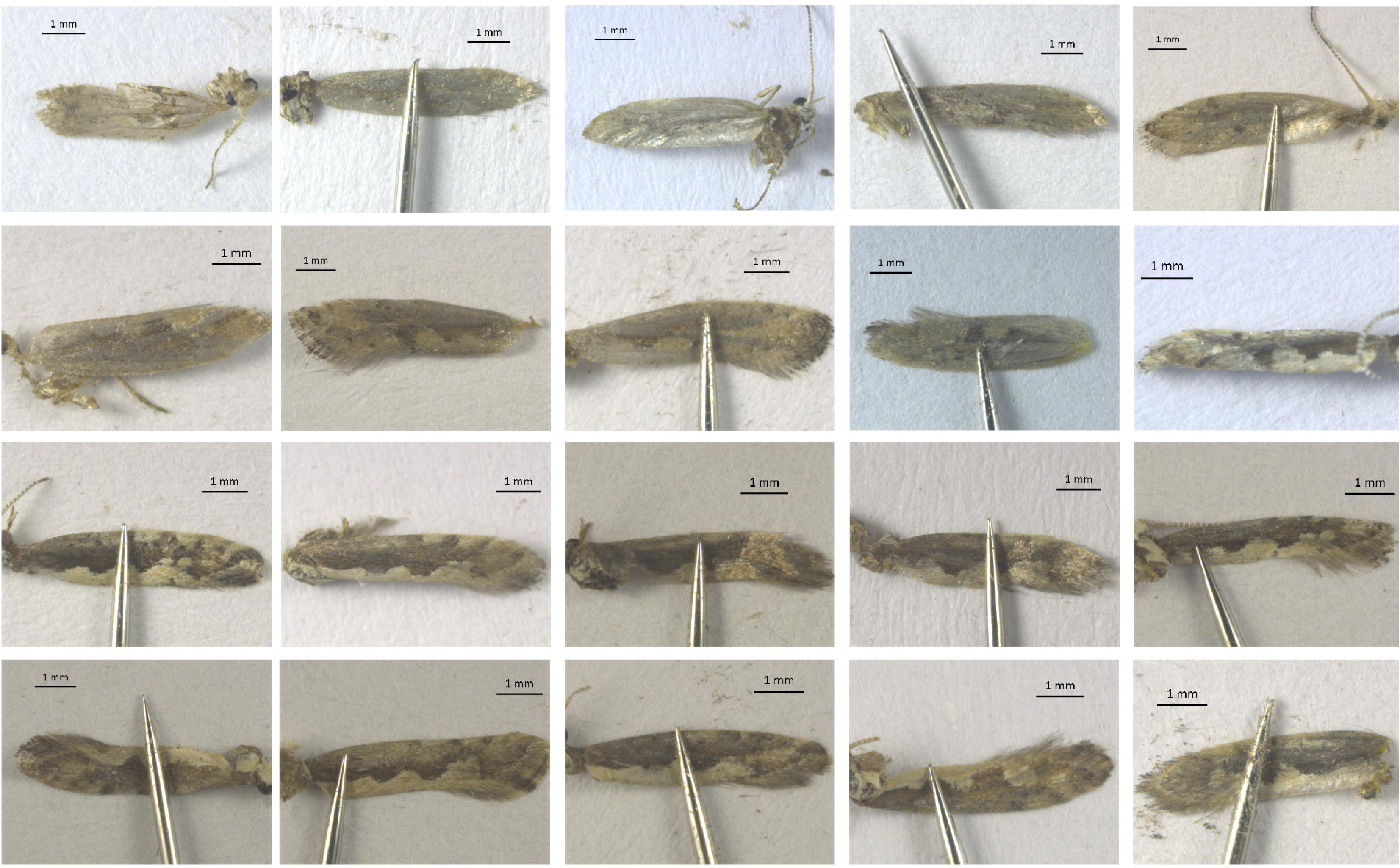
Expression levels of the *PxGP* gene after double-stranded RNA injection. Data are presented as mean ± SEM (n = 3); \*\*\**P* < 0.001, ds*GP* vs. ds*GFP* treatment group (independent samples t-test).

**Figure 10–figure supplement 2.**
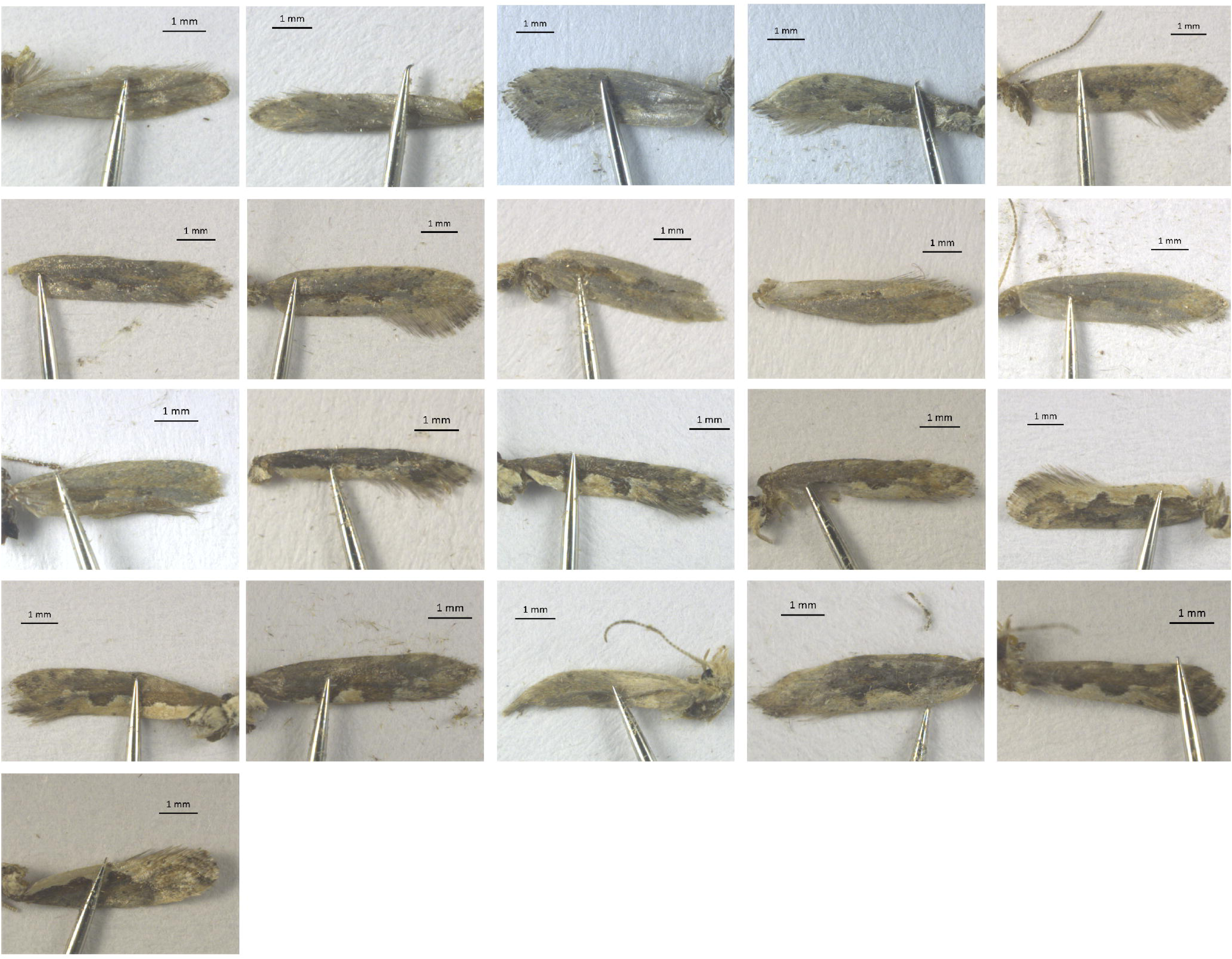
Total protein concentration in test insects subjected to glycogen phosphorylase activity determination, analysis of alternative glycogen-metabolizing enzymes, and fitness cost evaluation. Data are mean ± SEM (n=3). Significance vs. control: \*\**P* < 0.01 (two-way ANOVA with Sidak’s test).

**Figure 11–figure supplement 1.**
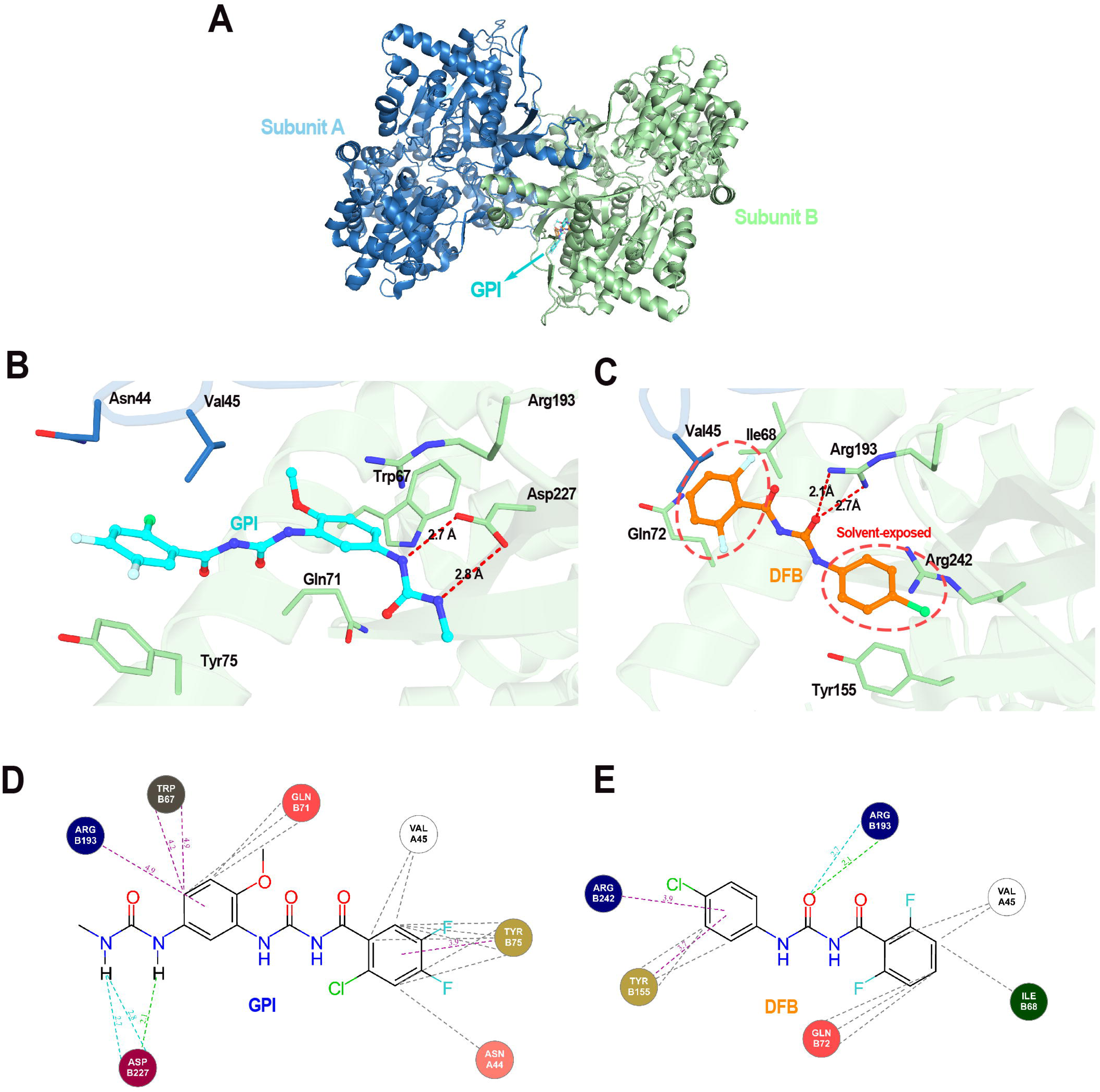
Wing phenotype of adult insects emerged from the ds*GFP*-treated group.

**Figure 11–figure supplement 2.**
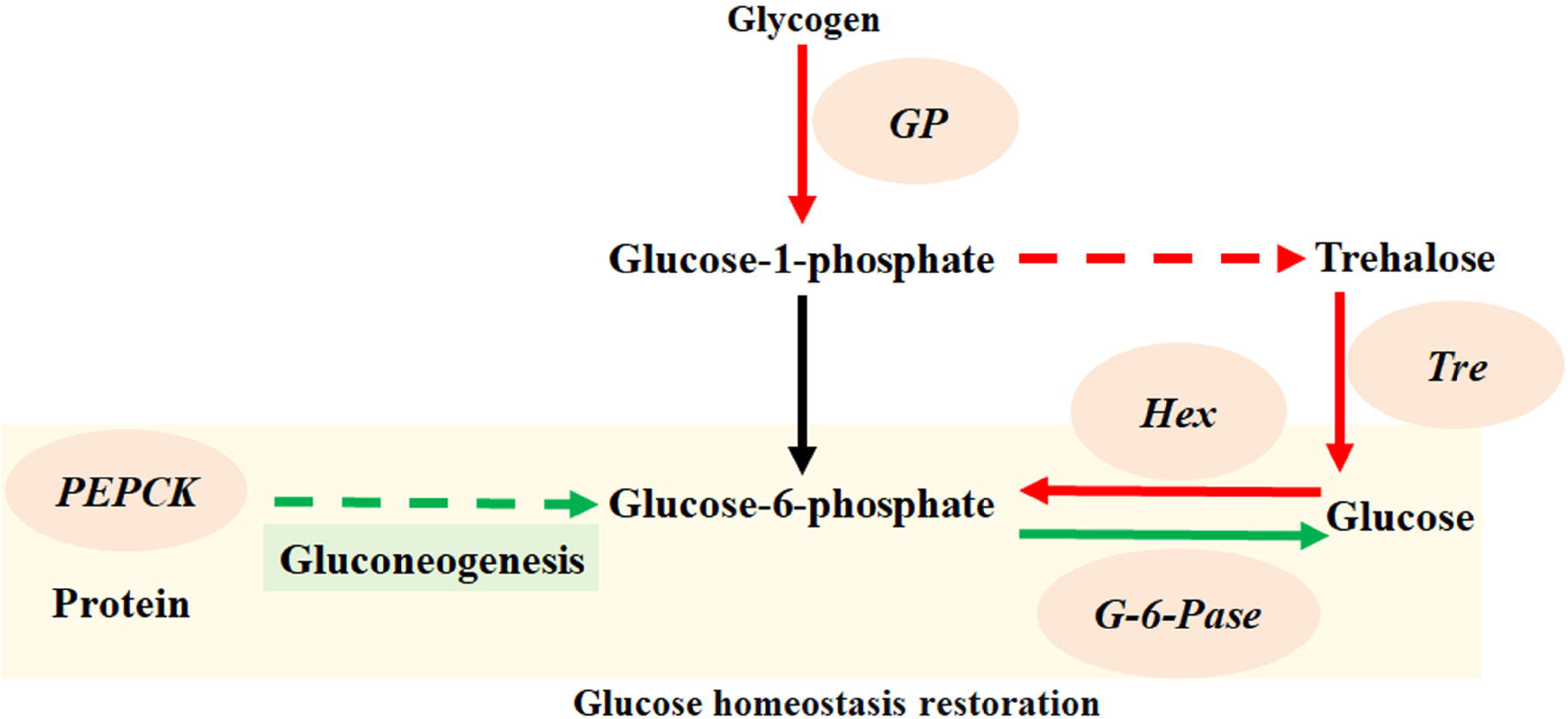
Wing phenotype of adult insects emerged from the ds*GP*-treated group.

## Additional information

### Data availability

All relevant data are within the paper and its supplementary files.

## Additional files

**Supplementary file 1.** Primer sequences used in this study.

**Supplementary file 2.** Toxicity of Diflubenzuron (DFB) against 3rd-instar *Plutella xylostella* larvae at 120 h.

**Source data 1.** Contains values used for data presentation of Figures 2-4, 6-11, and Figure 2–figure supplement 1, Figure 10–figure supplements 1 and 2.

