## Supplementary material for "Metabolic compensation via gluconeogenesis explains the non-essentiality of glycogen phosphorylase as an insecticidal target in *Plutella xylostella*": Primer sequences used in this study.

**Table S1.** Primer sequences used in this study.

| Primer Name | Sequence (5'-3') | Amplicon (bp) | Efficiency (%) | R² |
| --- | --- | --- | --- | --- |
| **RT-qPCR** |  |  |  |  |
| qPxGP-F | ACCCCAACGACCACTTCTTC | 170 | 107.86 | 0.986 |
| qPxGP-R | CGACCTTCTCAGGGAGGCTA |  |  |  |
| qPxTre-F | GCTCTACAACGACTCCAAG | 139 | 105.35 | 0.997 |
| qPxTre-R | CTGCGACACGAACTCCT |  |  |  |
| qPxHEX-F | CTGGGATTCACATTCAGTTT | 136 | 90.27 | 0.994 |
| qPxHEX-R | TGGCAATAGCGTCTTTCA |  |  |  |
| qPxpepck-F | TGTCAACTGGTTCCGCAAGAATG | 160 | 102.08 | 0.996 |
| qPxpepck-R | TGTGTTCAAGCAGCCGTCTCT |  |  |  |
| qG-6-P-F | GAGTTACTGCTGATTGTGATGTGGAT | 200 | 98.17 | 0.995 |
| qG-6-P-F | AGGGCTGGTGCTAGGAATGC |  |  |  |
| qPxGBE-F | TTCAACTTCAATACGACGCAGAG | 105 | 98.96 | 0.997 |
| qPxpepck-R | TCCGCCATATTCCTTGCTATCC |  |  |  |
| qPxα-Amylase-F | CTACGCTTGTTGTGCTGATAGG | 193 | 97.28 | 0.995 |
| QPxα-Amylase-R | TCTGCTGTAGATGACGCTGAG |  |  |  |
| qPRS13-F | TCAGGCTTATTCTCGTCG | 123 | 91.22 | 0.990 |
| qPRS13-R | GCTGTGCTGGATTCGTAC |  |  |  |
| **dsRNA Synthesis** |  |  |  |  |
| dsPxGP-F | GAATGTGACGGAGGTGAAGAAG |  |  |  |
| dsPxGP-R | ATGGCGAACACGATCTGAGT |  |  |  |
| dsT7PxGP-F | taatacgactcactatagggGAATGTGACGGAGGTGAAGAAG |  |  |  |
| dsT7PxGP-R | taatacgactcactatagggATGGCGAACACGATCTGAGT |  |  |  |
| dsGFP-F | GAGAAGAACTTTTCACTGCA |  |  |  |
| dsGFP-R | TGTTGATAATGGTCTGCTAG |  |  |  |
| dsT7GFP-F | taatacgactcactatagggGAGAAGAACTTTTCACTGCA |  |  |  |
| dsT7GFP-R | taatacgactcactatagggTGTTGATAATGGTCTGCTAG |  |  |  |
| **Protein expression** |  | |  |  |
| pFastbac-F | TATTCCGGATTATTCATACC | |  |  |
| pFastbac-R | ACAAATGTGGTATGGCTGA | |  |  |
| pUC/M13-F | CCCAGTCACGACGTTGTAAAACG | |  |  |
| pUC/M13-R | AGCGGATAACAATTTCACACAGG | |  |  |

Note: Lowercase represents the T7 promoter sequence: taatacgactcactataggg.

The qPCR amplification efficiency is represented by Efficiency (%).
