## Supplementary material for "Metabolic compensation via gluconeogenesis explains the non-essentiality of glycogen phosphorylase as an insecticidal target in *Plutella xylostella*": Toxicity of Diflubenzuron (DFB) against 3rd-instar Plutella xylostella larvae at 120 h.

**Table S2.** Toxicity of Diflubenzuron (DFB) against 3rd-instar *Plutella xylostella* larvae at 120 h.

| toxicity regression equation | LC_30_（mg/L） | LC_50_（mg/L） | χ^2^ | *P* |
| --- | --- | --- | --- | --- |
|  | (95% CI) | (95% CI) |  |  |
| y=-5.927+2.511x | 141.900  （67.893-206.576） | 229.537  （145.287-331.784） | 11.946 | 0.683 |
